# Multi-omics Differential Inference for Functional Interpretation (MoDIFI): A Statistical Framework to Prioritize Cell Lines for Neurodevelopmental Variants

**DOI:** 10.64898/2026.01.29.702065

**Authors:** VR Arvinden, Grace Tzun-Wen Shaw, Juana Manuel, Timothy L. Mosbruger, Hillary Heins, Jeffrey K. Ng, Haedong Kim, Tristan J. Hayeck, Tychele N. Turner

## Abstract

Noncoding variants contribute to neurodevelopmental disorders (NDDs), but their regulatory effects are often cell-type specific, making it difficult to choose an *in vitro* model for high-throughput assays such as massively parallel reporter assays. We asked: given a set of noncoding variants, which cell line and regulatory regions are most likely to reveal measurable allele-specific effects? We generated matched multiomics profiles across commonly used NDD *in vitro* models: human neuronal lines (i.e., IMR-32, SH-SY5Y, SK-N-SH), mouse neuronal lines (i.e., HT-22, Neuro-2a), and a non-neuronal line (i.e., HEK-293), using RNA-seq, ATAC-seq, and Hi-C under consistent conditions. To integrate these orthogonal data types, we developed MoDIFI (Multi-omics Differential Inference for Functional Interpretation), a Bayesian framework that quantifies cell-line-specific regulatory activity by computing posterior inclusion probabilities (PIPs) for differential gene-loop interactions. MoDIFI identifies regulatory regions supported by coordinated 3D contacts, accessibility, and transcriptional output, producing cell-line-resolved regulatory maps that highlight both shared synaptic programs and context-dependent mechanisms. These results provide a practical strategy for prioritizing the most informative cell lines and candidate regulatory elements for targeted functional testing of NDD-relevant noncoding variation.

## INTRODUCTION

Large-scale genomic studies of neurodevelopmental disorders (NDDs) have identified more than 300 genes as important contributors to these phenotypes.^1,2^ Increasing use of genomic sequencing to understand and diagnose NDDs has led to the identification of relevant variants, often in protein-coding regions. Some studies have reported an enrichment of variants in certain noncoding regions (e.g., promoters, enhancers),^3–6^ though the broader functional consequences of these noncoding signals are still not well characterized. Unlike protein-altering variants, noncoding variants have the potential to modify the expression of their target gene via various epigenetic and higher-level chromatin organization. Various available methods including Deep Mutational Scanning (DMS), Massively Parallel Reporter Assays (MPRA),^7,8^ and Multiplexed Assays of Variant Effects (MAVE)^9–14^ have enabled us to assess the functional role of coding and noncoding regions at scale. One of the primary challenges in these studies is the choice of the *in vitro* model used to perform these assays. This is particularly important for noncoding variants as different cell lines or cell types have distinct regulatory landscapes. ^7,89–14^ Use of multiomics data and scalable experimental systems creates an opportunity to link regulatory variation to functional outcomes, but data and methods to fully leverage these resources remain limited at this time.

As a result, prioritizing specific candidate regulatory elements specific to a given cell line or tissue while leveraging multiomics evidence is critical to identifying the most informative targets for such assays. Therefore, methods that can isolate targeted regulatory regions in relevant tissues or cell lines are needed as an initial step to guide noncoding variant highthroughput *in vitro* experiments. Existing methods link multi-omics data (e.g., Activity-by-Contact (ABC) model^15^) and have been widely used in conjunction with CRISPR-based screens to isolate key regulatory elements. Methods such as SNP-to-gene (S2G)^16^ mapping and related approaches have been effectively used to combine disease-related variants with target genes and disease linking multi-omics data. Such methods have proven to be powerful at leveraging experimental information into inference. However, these strategies typically rely on testing a given tissue or cell-type and do not address how regulatory activity varies across cell lines or how to combine orthogonal evidence to improve inference beyond linking them. This is partly because, until recently, there has been a lack of comprehensive, well-curated multi-omic datasets spanning multiple cell lines, with each cell line profiled for 3D interactions (e.g., Hi-C), chromatin accessibility and epigenetic state (e.g., ATAC-seq, DNA methylation), and gene expression (e.g., RNA-seq). While many studies (most notably ENCODE,^17,18^ which remains the largest public resource for these types of multi-omic datasets) include some of these components for individual cell lines, few provide all three layers for the same line. Given the strong cell-type specificity of noncoding regulatory elements, choosing the right cell lines is essential.

The central question driving this work was: given a large set of noncoding variants of interest, which cell line and regulatory region would maximize our ability to detect functional differences between reference and alternate alleles? To address this, we generated a comprehensive multi-omics dataset across cell lines commonly used in neurodevelopmental disorder (NDD) research. These included neuronal models (human SH-SY5Y, SK-N-SH, IMR-32; mouse HT-22, Neuro-2a) and a nonneuronal model (human HEK-293). For each cell line, we performed RNA-seq, ATAC-seq, and Hi-C experiments under consistent conditions. This experimental framework ensured uniformity across cell lines and enabled direct comparisons of regulatory landscapes. Furthermore, beyond generating integrated multi-omics profiles across these cell lines, we also introduce a Bayesian framework to improve inference of differential regulatory activity. We present our framework Multiomics Differential Inference for Functional Interpretation (MoDIFI, i.e. “modify”), which quantifies differential regulatory activity at gene-loop pairs by integrating evidence from gene expression, chromatin accessibility, and 3D genome interaction across cell lines. MoDIFI calculates posterior inclusion probabilities (PIPs) for each gene-loop pair, providing a probabilistic measure of whether an interaction is functionally differential in one cell line versus others. This framework enables identification of cell-line-specific regulatory regions that can guide targeted experimental follow-up that could be prioritized for MPRA or similar saturation mutagenesis approaches.

## RESULTS

### Multi-Omic Data Generation for MoDIFI

To develop the model (Figure 1) for assessing cell type suitability in high throughput regulatory region screening, we first generated a multi-omics dataset capturing transcriptomic, chromatin accessibility, and 3D genome organization across six cell lines: human neuroblastoma-derived neuronal cell lines (SH-SY5Y, SK-N-SH, IMR-32), mouse neuronal (HT-22, Neuro-2a), and human non-neuronal baseline (HEK-293), which are derived from human embryonic kidney tissue. All cell lines were cultured under standardized conditions and profiled using complementary assays to capture distinct layers of gene regulation. Specifically, we performed RNA-seq to quantify transcript abundance, ATAC-seq to map open chromatin regions, and Hi-C to characterize 3D genome architecture. Each dataset was processed using the appropriate ENCODE pipeline for the respective assay, followed by additional post-processing and filtering as required for downstream analyses. These experiments were conducted uniformly across all cell lines to enable direct comparisons of regulatory landscapes. Summary results from ATAC-seq open chromatin regions, Hi-C loop calling for 3D interactions, and RNA-seq expression across neuronal cell lines and the comparator cell line HEK-293 show relatively low overlap across all annotations (Figure **2**2A, **Error! Reference source not found**., and see Supplemental Data).

**Figure 1.**
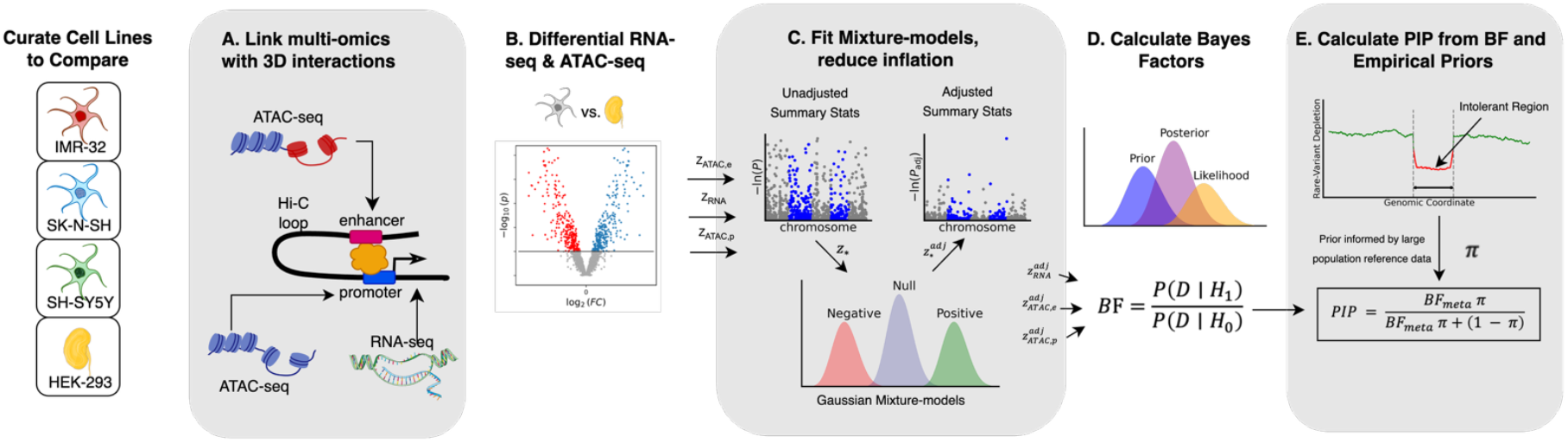
MoDIFI: A Bayesian Framework for Integrating Multi Omic Evidence Across Cell Lines. Hi-C, ATAC-seq, and RNA-seq experiments provide cell line specific regulatory, chromatin, and transcriptional signatures. These modalities are linked through Hi-C loops, differential expression and accessibility are quantified and corrected using mixture model calibration, and combined through Bayes factors. Empirical priors derived from a large population genetics reference cohort calibrate the final posterior inclusion probabilities, yielding cell type specific regulatory interaction estimates.

### Overview of Bayesian Framework

We introduce MoDIFI, a Bayesian framework that integrates cross cell line evidence from gene expression, chromatin accessibility, and 3D genome interaction to quantify relative differential gene-loop activity in a target cell line compared to all others or a designated baseline (here, HEK-293 serves as the baseline for neuronal cell lines since it is a non-neuronal cell line). The workflow consists of four steps. First, Hi-C data are used to define interaction loops, and summary statistics for RNA-seq and ATAC-seq are obtained using DESeq2^19^ to estimate differential expression and differential chromatin accessibility, followed by adjustment for bias and inflation via a mixture model to ensure interpretability across modalities (**Error! Reference source not found**. and **Error! Reference source not found**.).^20^ Second, component Bayes factors are computed from these adjusted z-scores for RNA-seq and ATAC-seq evidence, with ATAC partitioned into promoter-proximal and distal regulatory ends of Hi-C loops, and combined within each cell line using a weighted log-BF scheme to stabilize inference and avoid inflation from naive multiplication. Third, for comparisons involving multiple cell lines, evidence is aggregated by computing a weighted average of log Bayes factors across comparators to obtain a meta-level BF for each loop–gene pair. Finally, the aggregated Bayes factor and priors are used to calculate a posterior inclusion probability (PIP) using biologically informed priors. We adopt an empirical Bayes strategy that leverages orthogonal evidence of evolutionary constraint (derived from Gnocchi scan statistics^21^ on Hi-C loop regions) to prioritize regions under negative selection, improving inference over uninformative flat priors. Full details, including Bayes factor derivations, weighting schemes, and prior estimation, are provided in the Supplementary Methods.

### Gene Family Enrichment Shows Key Gene Families with Differential Omics Results Across Cell Line

Using the super gene families defined by the Human Genome Organization (HUGO) Gene Nomenclature Committee (HGNC),^22^ we tested for relative enrichment across the different omics data for each cell line. There were a total of 38,854 unique human genes with 16,990 orthologues to a set of 31,268 mouse genes. First, pairwise analysis was conducted across each cell line, utilizing RNA-seq results for differential expression and ATAC-seq results linked with Hi-C loops to assess differential chromatin accessibility and interactions. The analyses aimed to assess whether there is an enrichment or increased frequency of genes in each super gene family (relative to the entire genome) that show differential expression or differential open chromatin regions (with interactions) falling within the top 10%.

For Neuro-2a and HT-22, ATAC-seq regions overlapping gene promoters and Hi-C loops are shown in Figure 2. Gene set enrichment analysis identified the proportion of genes within each superfamily in the top 10% of differentiation (Figure **2**2B). Then an exact test was performed to assess the likelihood of this enrichment in fold change occurring by chance within each superfamily for each cell line.

**Figure 2.**
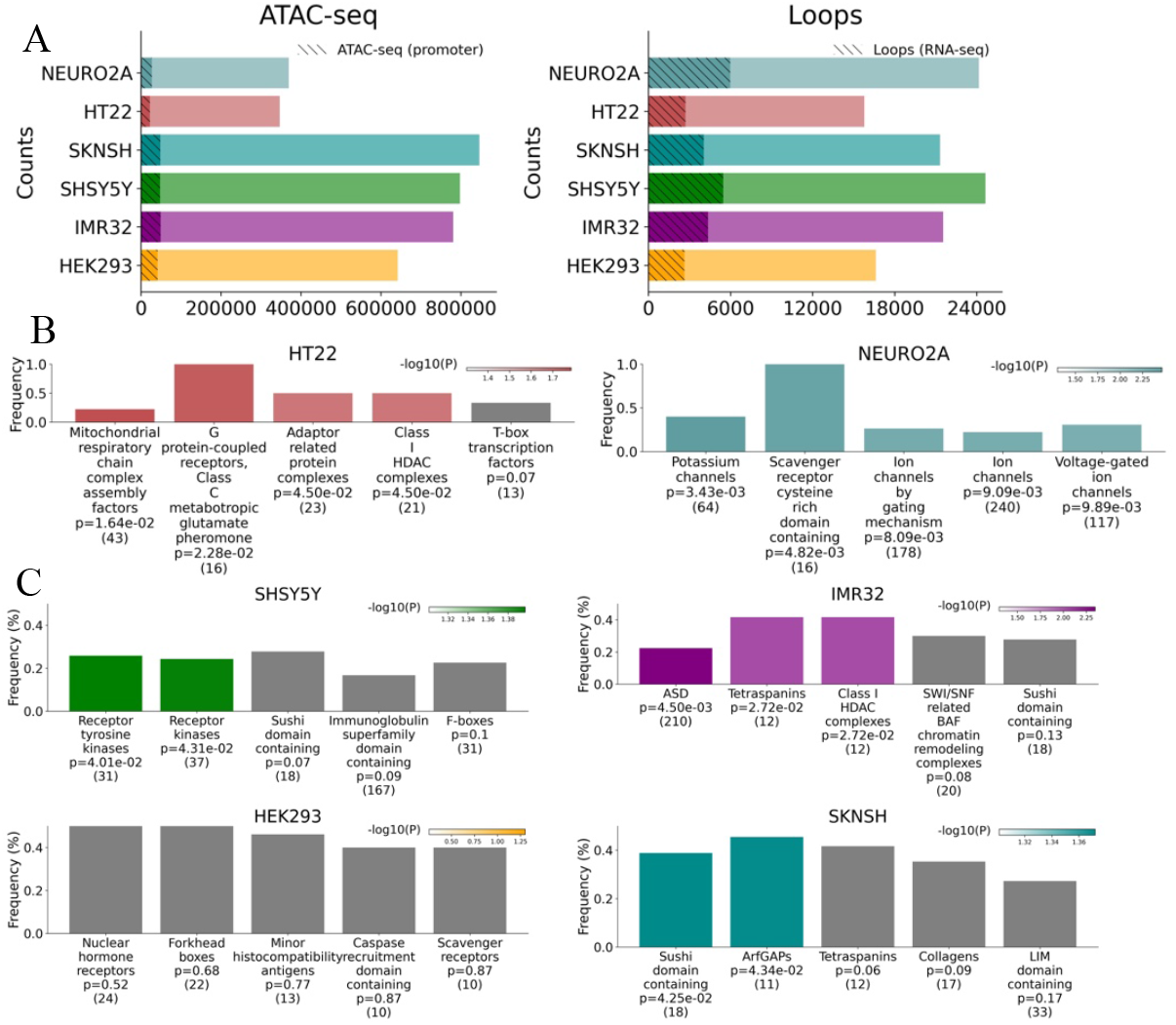
Functional data across cell lines and HGNC super gene families. For each cell line A.) counts of ATAC-seq peaks and Hi-C loops across the different cell lines. Then there is sub-bar plots shaded indicating those ATAC-seq regions that also hit a promoter and those loops that correspond to RNA-seq expression data as well. Functional gene family proportions across HGNC super gene families for B.) mice and C.) humans cell lines. For Neuro-2a and HT-22 enrichment of ATAC-seq regions hitting gene promoter and with Hi-C loops are shown, while for SK-N-SH, IMR-32, SH-SY5Y, and HEK-293 cell type-specific differential multiomics activity and regulatory activation PIP enrichment are shown for HGNC functional gene families. The top ten super gene families with the highest frequencies of highly differentiated genes (top 10%) per cell line are shown, ranked by significance. Darker colors indicate lower p-values, highlighting superfamilies most enriched for regulatory function. The number of genes in each supergene family is shown in parentheses.

For the human cell lines examined in this study, we quantified, for each HGNC superfamily, the proportion of genes that were differentially expressed and ranked in the top 10% (Supplemental Figure 3). This data was then used to conduct an exact test to assess the probability of observing this level of relative enrichment by chance. In IMR-32, we observed significant enrichment for ion channels, particularly potassium and voltage-gated channels, with some families comprising over 40% of the genes in the top percentage. Using an independently curated autism spectrum disorder (ASD)-related gene set from the literature, IMR-32 also showed marginally significant enrichment at an alpha level of 5% for differentially expressed genes (*p* = 0.027). Consistently, genes containing *de novo variants* (DNVs) identified in individuals with ASD were significantly enriched as well (*p* = 1.5 × 10^−4^).

Next, we performed enrichment tests on protein-coding genes whose promoters were substantially covered by ATAC-seq peaks (>50% of the promoter region) and that were also anchored by Hi-C loops, capturing loci that are both accessible and potentially engaged in long-range regulatory interactions (Supplemental Figure 4). Focusing on IMR-32, we again observed significant enrichment for ion channel families, especially potassium, and voltage-gated channels, with several HGNC superfamilies overlapping those identified in the RNA expression-based analysis. Moreover, genes previously implicated in ASD met nominal significance (*p* = 0.02).

Last, across the human derived cell lines we investigated enrichment differential multi-omics activity and regulatory activation, as measured by our differential activity and contact framework, MoDIFI (see Methods).

In human brain cell lines, we quantified the percentage of PIP that fell within the top 5% (i.e., PIP ≥ 95%), using HEK-293 cells as a reference (Figure **2**2C). In the case of a gene promoter involved in multiple loops or that is within a loop that is linked to multiple ATAC-seq regions, a median PIP of all interactions was used to represent the gene. Similarly, exact tests were performed to see if the percentage of genes within a given super gene family, *p*_*genes*_, with median PIP in the top 5% was elevated for that group versus all others. Distinct gene family enrichments were observed across neuroblastoma-derived neuronal cell lines. IMR-32 showed significant enrichment for ASD-associated genes (*p*_*genes*_ = 22.38%,, *p* = 4.5 × 10^−3^), tetraspanins (*p*_*genes*_ = 41.67%, *p* = 2.72 × 10^−2^) and class I HDAC complexes (*p*_*genes*_ = 41.67%, *p* = 2.72 × 10^−2^). SH-SY5Y cells exhibited enrichment for receptor tyrosine kinases (RTKs) (*p*_*genes*_ = 25.81%, *p* = 4.01 × 10^−2^) and receptor kinases (*p*_*genes*_ = 24.32%, *p* = 4.31 × 10^−2^). SK-N-SH cells showed enrichment for sushi domain–containing proteins (*p*_*genes*_ = 38.89%, *p* = 4.25 × 10^−2^) and ArfGAP family genes (*p*_*genes*_ = 45.45%, *p* = 4.34 × 10^−2^).

The observed gene family enrichment patterns are consistent with known functional distinctions among neuroblastoma-derived neuronal models. In IMR-32, enrichment of ASD-associated genes and class I HDAC complexes aligns with evidence that epigenetic mechanisms, including histone deacetylase activity, play important roles in early neurodevelopment^23,24^ and ASD.^25,26^ Class I HDACs, such as HDAC3, regulate chromatin accessibility during neuronal differentiation and contribute to neurodevelopmental gene regulation through modulation of transcriptional programs.^27^ Tetraspanins, which modulate membrane receptor organization and neurite development^28^ support cellular programs involved in synaptic structuring and receptor signaling that are frequently implicated in ASD-relevant pathways.^29,30^ Their selective enrichment in IMR-32, but not in SK-N-SH or SH-SY5Y, suggests cell line-specific differences in neurodevelopmental or connectivity-related regulatory programs. In contrast, enrichment of sushi domain-containing proteins and ArfGAP family genes in SK-N-SH highlights molecular mechanisms related to membrane dynamics, vesicular trafficking, and synaptic organization, suggesting an emphasis on synaptic maintenance and structural regulation.^31–34^ SH-SY5Y cells showed enrichment for receptor tyrosine kinases and receptor kinases, consistent with their established use as a model for neuronal signal transduction, differentiation, and neurotrophic factor responsiveness. Notably, several RTKs implicated in ASD, such as MET, play critical roles in early neuronal growth and circuit formation.^35,36^ but continue to influence synaptic signaling and refinement at later stages, suggesting that SH-SY5Y captures sustained RTK-mediated pathways with relevance across neurodevelopmental timelines.

### Integration of PGC Association Study and Multi-Omics Data: Exploring Trait-Associated Variants and Differential Functional Evidence

Variants associated with psychiatric and neurodevelopmental disorders from the PGC are expected to occur more frequently in regions with highly differentiated functional activity. This suggests that these variants may show a trend of increasing variant-trait association significance with higher MoDIFI PIP values in cell lines that correspond to the biological function of the trait. To test this, we utilized a range of PGC traits with available association study data such as attention deficit hyperactivity disorder (ADHD),^37^ Bipolar Disorder (BIP),^38^ Anxiety Disorder (ANX),^39^ Major Depressive Disorder (MDD),^40^ eating disorders (ED),^41^ obsessive compulsive disorder and Tourette Syndrome (OCD-TS),^42^ Post Traumatic Stress Disorder (PTSD),^43^ schizophrenia (SCZ), and Substance Use Disorders (SUD) to check if significantly associated variants are enriched in high MoDIFI PIP bins. For BIP and PTSD, analyses were further stratified by available genetic ancestries (**Error! Reference source not found**.5). For BIP, these included African (AFR), East Asian (EAS), European (EUR), and a multi-ancestry meta-analysis (Multi), whereas for PTSD, stratification included African (AFR), Hispanic/Native American (AMR), European (EUR), and a multi-ancestry meta-analysis (Multi). Additionally, we investigated associations across results from a recent large cross-disorder genomic study^44^ and applied latent factor modeling to genome-wide association data across 14 psychiatric disorders, identifying five major factors cross disorder dimensions: F1 Compulsive, F2 Schizophrenia/Bipolar, F3 Neurodevelopmental, F4 Internalizing and F5 Substance Use Disorders. F1, a Compulsive disorders factor defined by Anorexia nervosa (AN), Obsessive-compulsive disorder (OCD), and, more weakly, Tourette’s syndrome (TS) and Anxiety disorders (ANX); F2, a Schizophrenia/Bipolar (SB) factor defined by Schizophrenia (SCZ) and Bipolar disorder (BIP); F3, a Neurodevelopmental factor defined by ASD, Attentiondeficit/hyperactivity disorder (ADHD), and, more weakly, TS; F4, an Internalizing disorders factor defined by Posttraumatic stress disorder (PTSD), Major depression (MD), and ANX; F5, a Substance Use Disorders (SUD) factor defined by Opioid use disorder (OUD), Cannabis use disorder (CUD), Alcohol use disorder (AUD), Nicotine dependence (NIC), and, to a lesser extent, ADHD (**Error! Reference source not found**.6).

Since individual chromatin loops could contain multiple interaction combinations, the maximum MoDIFI PIP was used to represent each loop. PGC variants were then mapped to these unique loops, and when a variant mapped to multiple loops, the mean MoDIFI PIP across all mapped loops was assigned to that variant. First, for each cell line, we evaluated whether SNPs with −*log*_10_(*p*) values at or above a given threshold were associated with higher odds of being strongly functionally active, defined as the top 5% of MoDIFI PIP values, compared to those below this threshold. Exact tests were similarly performed looking at SNPs with −*log*_10_(*p*) values at or above a given threshold versus being strongly functionally active (top 5% PIP) versus not. Clear trends emerged across specific cell lines for certain traits (Figure 3 and Figure 4A). To quantify these patterns and determine whether the OR of variants in the top 5% versus bottom MoDIFI PIP bins shows associations with variant-trait significance levels (−log_10_(p) bins), we applied the Greenland–Longnecker^45^ trend test, fit using significance bin (−*log*_10_(*P*) > {2,3,4,5}). The upper threshold was chosen at (10^−5^) as a bit below genome-wide significance to capture variants that might otherwise be missed or lie near the significance boundary, where additional evidence could strengthen association, and because OR confidence intervals beyond that level started to show considerably higher variability, likely due to small group size. The Greenland-Longnecker method provides a robust approach for estimating associations while accounting for confidence intervals, commonly applied in meta-analysis.

**Figure 3.**
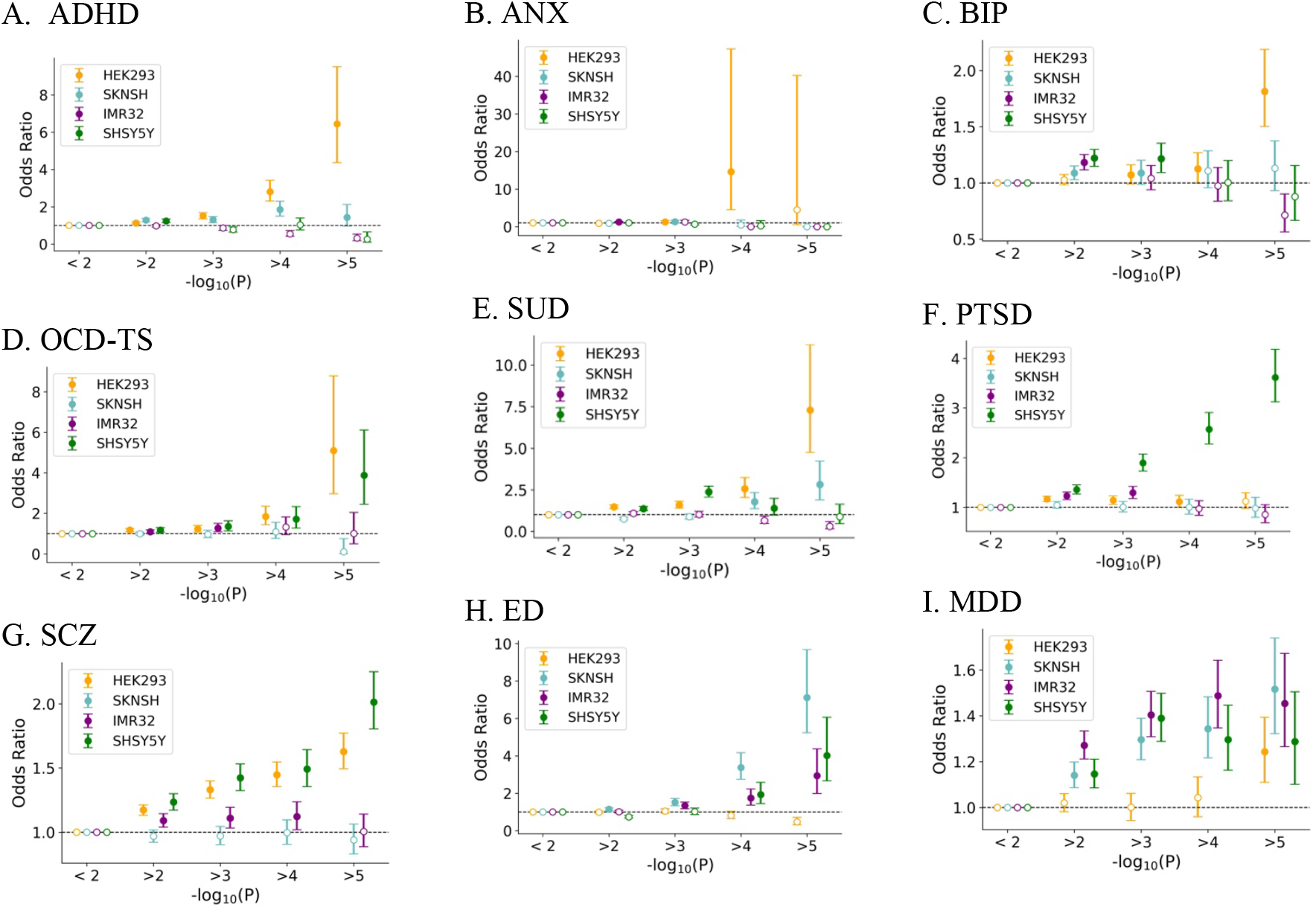
Enrichment of psychiatric disorder–associated variants in high MoDIFI PIP regulatory regions across cell lines. For each PGC trait, panels display the odds ratio of SNPs overlapping the top 95th percentile of posterior inclusion probability (PIP) scores versus those below the 95th percentile across four cell lines (HEK-293, SK‐N‐SH, IMR‐32, and SH‐SY5Y). SNPs are binned by increasing GWAS significance (−log_10_(p) thresholds), and points represent odds ratios with 95% confidence intervals error bars. Horizontal dashed lines denote no enrichment (OR = 1). For all plots, solid dots indicate significant OR at the 5% alpha level whereas hollow ones are not significant.

**Figure 4.**
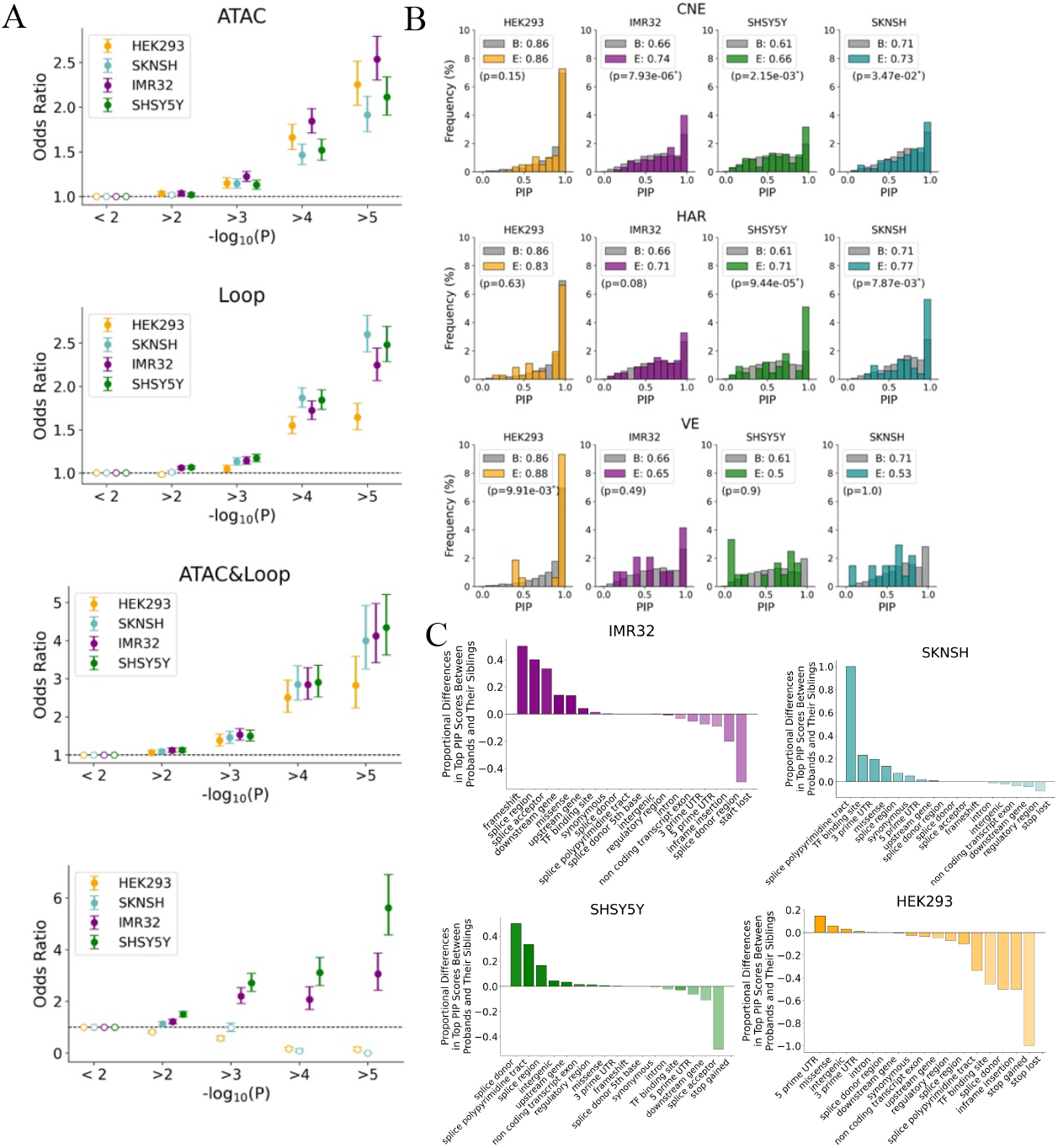
MoDIFI PIP functional omics trends across ASD‐associated GWAS variants, DNVs, and linked enhancer regions. Investigating A.) trends between functional data and −*log*_10_(*p*) values from the PGC ASD GWAS study are shown. In the lowest pane, the association of SNP odds ratios with the top 5% of MoDIFI scores (i.e., MoDIFI ≥95%) across different cell lines, using −*log*_10_(*p*) values from the PGC ASD study (similar to Figure 3 for other traits). For all plots, solid dots indicate significant OR at the 5% alpha level, whereas hollow ones are not significant. In B.) the distribution of frequency of PIP values for loops connecting to enhancer7 is plotted in comparison to the PIP PIP values for all loops, excluding those already identified as connecting to enhancers previously identified as enhancer‐associated. Enhancers are derived from HARs, VEs, and CNEs linked to ASD.^47^ For C.) we examine functional annotations for DNVs with high MoDIFI scores in probands versus siblings in the SFARI cohort. Bar plots display the difference in proportions of DNVs ranking in the top 5% of MoDIFI scores between probands and siblings, where DNVs are grouped by VEP consequence categories.

For each cell line and corresponding trait with a significant Greenland-Longnecker trend, results are summarized below. The association estimator from the Greenland-Longnecker test is reported as β, with its corresponding p-value (*p*_*GL*_) *β, p*_*GL*_, −*log*_10_(*p*)]. Odds ratios at the highest GWAS significance bin (−*log*_10_(*P*) > 5) are reported as *OR*_*high*_, with their p-value (*p*_high_). Looking by cell line, significant results were observed in SH-SY5Y for PTSD (*β*_*GL*_ = 0.297, *p*_*GL*_ = 0.0016; *OR*_*high*_ = 3.08, *p*_*high*_ = 8.48 × 10^−46^) and SCZ (*β*_*GL*_ = 0.113, *p*_*GL*_ = 0.0255; *OR*_*high*_ = 1.8112, *p*_*high*_ = 6.11 × 10^−26^). Then for SK-N-SH for ED (*β*_*GL*_ = 0.523, *p*_*GL*_ = 0.038; *OR*_*high*_ = 7.118, *p*_*high*_ = 1.39 × 10^−34^), CDG F1 Compulsive Disorders factor (*β*_*GL*_ = 0.421, *p*_*GL*_ = 0.051; *OR*_*high*_ = 6.149, *p*_*high*_ = 1.46 × 10^−15^), MDD (*β*_*GL*_ = 0.092, *p*_*GL*_ = 0.02; *OR*_*high*_ = 1.52, *p*_*high*_ = 5.94 × 10^−9^), and for CDG F4 Internalizing Disorders factor (*β*_*GL*_ = 0.092, *p*_*GL*_ = 0.061 ;*OR*_*high*_ = 1.51, *p*_*high*_ = 1.23 × 10^−5^) significant results were observed in. In contrast, HEK-293 showed its strongest enrichment for ADHD (*β*_*GL*_ = 0.442, *p*_*GL*_ = 0.037; *OR*_*high*_ = 6.44, *p*_*high*_ = 7.08 × 10^−29^) and the cross-disorder CDG F2 schizophrenia/bipolar–related factor (*β*_*GL*_ = 0.23, *p*_*GL*_ = 0.004; *OR*_*high*_ = 2.096, *p*_*high*_ = 1.40 × 10^−20^). In addition, significant enrichment was observed for the CDG F3 neurodevelopmental disorders factor (*β*_*GL*_ = 0.890, *p*_*GL*_ = 0.0292), although the effect at the most stringent GWAS threshold was attenuated (*OR*_*high*_ = 3.60, *p*_*high*_ = 0.248).

Next, focusing on ASD^46^ from the PGC in more detail as an illustrative example (Figure 4). ASD was chosen in part because we have additional follow-up data available, including DNVs across a large set of parent-child sequenced trios for both probands and unaffected siblings. This allowed us to examine individual multi-omics components in greater detail and when applying our MoDIFI framework across cell lines to assess its consistency and potential utility for this trait. When examining the individual contributions of the different functional evidence, a general trend indicated that, typically, the more significant the p-value, the higher the odds ratio for the variant being in both a loop and an ATAC region (Figure 4A). When looking across MoDIFI OR for ASD results demonstrated a significant trend (for SH-SY5Y (*β*_*GL*_ = 0.429, *p*_*GL*_ = 0.020; *OR*_*high*_ = 5.61, *p*_*high*_ = 4.38^−49^) (Figure 4A bottom pane), similar to what was previously seen with for ED, SCZ, and PTSD along with latent factors F1 and F2.

### Regulatory Activation of Enhancers related to ASD Risk

Recent work has shown that rare variation in noncoding regions with evolutionary signatures contributes to ASD risk,^47^ looking specifically at enhancers derived from Human Accelerated Regions (HARs)^48^, VISTA Enhancers (VEs), ^49^ and Conserved Neural Enhancers (CNEs).^47,50^ To further investigate this in the context of the functional experiments performed in this project, we looked to see if there was an elevation of cell type-specific differential multi-omics activity and regulatory activation linking to our data. The distribution of frequency of median multi-omics differential PIP for loops connecting to enhancers are plotted in comparison to the MoDIFI scores for all loops, excluding those already identified as connecting to enhancers (Figure 4**Error! Reference source not found.Figure** **2**B). Wilcoxon rank-sum tests comparing enhancer-associated and background loops revealed significant rightward shifts for loops connecting to CNEs in IMR-32 (*p* = 7.93 × 10^−6^; median shift +0.103, median 0.788, IQR = 0.569 − 0.955 vs. background *median* = 0.685, IQR=0.470 − 0.895), SH-SY5Y (*p* = 2.15 × 10^−3^; median shift +0.058, *median* = 0.681, IQR = 0.462 − 0.929 vs. *median* = 0.623, IQR = 0.408 − 0.830), and SK-N-SH (*p* = 3.47 × 10^−2^; median shift +0.052, *median* = 0.793, IQR = 0.577 − 0.947 vs. 0.741, IQR = 0.561 − 0.906), as well as for HARs in SH-SY5Y (*p* = 9.44 × 10^−5^; median shift +0.102, *median* = 0.725, IQR = 0.466 − 1.000 vs. 0.623, IQR 0.408 − 0.830) and SK-N-SH (*p* = 7.87 × 10^−3^; median shift +0.069, *median* = 0.810, IQR = 0.611 − 0.997 vs. 0.741, IQR = 0.561 − 0.906). In contrast, IMR-32 showed weaker enrichment for HAR-associated loops (*p* = 0.08), and VISTA enhancer (VE)-associated loops displayed little or no significant enrichment across neuronal cell types. Notably, in cell lines other than IMR-32, multi-omics differential PIP are typically higher among loops connecting to HARs and CNEs, showing statistically significant shifts from performing Wilcoxon-tests across the two distributions.

### Linking Omics Data and Functional Consequences Comparing DNVs in ASD Probands and Control Sibling

DNVs play a major role in ASD,^6,51–55^ hence we examined DNVs from large ASD cohorts comprising probands and unaffected sibling controls. Focusing on the proportion of DNVs that fell within functional omics data, either Hi-C interaction data or ATAC-seq open chromatin regions. We examined if functional annotations had consistent discrepancies between probands and siblings. We then stratified the variants into two groups: those found only in probands and those found only in siblings. For each group (probands and siblings) we linked the variants to the omics data from each cell line including Hi-C interaction data, ATAC-seq open chromatin regions, and both Hi-C interactions and open chromatin regions. Variants from proband and sibling groups were then classified into two categories based on MoDIFI scores (MoDIFI ≥ 95% and MoDIFI < 95%). We subsequently assessed whether probands exhibited a higher proportion of high-MoDIFI variants compared to siblings. Additionally, we associated the variants with high PIP scores, in the top 5% (i.e., MoDIFI ≥ 95%), to assess any differences in combined functional consequences between probands and siblings. This comprehensive approach allowed us to evaluate how these multi-omics data might reveal distinct patterns related to the DNVs in ASD probands versus control siblings.

Next, we compared by testing the relative proportions of variants associated with each functional annotation between probands and siblings. The significant results from either Fisher’s exact test or the proportion test are presented here. First, we examined the proportion of DNVs located within a Hi-C loop (Supplemental Table 1) and found significant differences between probands and siblings in the IMR-32 cell line for stop gain and intronic variants. In the SH-SY5Y cell line, significant differences were observed between probands and siblings for stop gain, splice region, and stop lost variants. For the SK-N-SH cell line, significant differences between probands and siblings were noted for splice donor 5th base variants. No significant results were found within HEK-293, which is possible because this was a baseline cell line used for comparison where differences were not necessarily expected.

We next examined ATAC-seq open chromatin regions to investigate the differences in functional annotations of DNVs, comparing those found within these regions (as compared to variants located outside the ATAC-seq open chromatin regions) in ASD probands to those in control siblings (Supplemental Table 2). We found significant differences in the HEK-293 cell line for TFBS, non-coding transcript variants, and splice donor region variants between probands and siblings. In the IMR-32 cell line for synonymous and splice acceptor variants showed significant difference between probands and siblings. In the SH-SY5Y cell line, significant differences were observed for only intergenic variants and lastly for the SK-N-SH cell line, significant differences were noted for start lost variants. Next, we examined overlaps between Hi-C interactions and open chromatin regions jointly. The proportion of variants with high PIP, ≥95%, was calculated for each VEP functional consequence category across probands and siblings to assess significant differences (Figure 4**Error! Reference source not found**.C and **Error! Reference source not found**. Table 2). Across neuronal cell lines, probands consistently showed a higher proportion of DNVs in the top 5% MoDIFI (i.e., MoDIFI ≥ 95%) category compared to siblings, with significant enrichments observed across multiple variant consequences (Figure 4C and Table 2). In IMR-32, both downstream gene variants (*p*_*diff*_ = 13.95%; p = 0.0076) and missense variants (*p*_*diff*_ = 13.60%; p = 0.0244) were significantly enriched in probands. In SH-SY5Y, intergenic variants showed a modest but significant enrichment by proportion testing (*p*_*diff*_ = 4.47%; p = 0.0495), and splice polypyrimidine tract variants exhibited a pronounced difference (*p*_*diff*_ = 33.33%; p = 0.0484), though with limited sample sizes. In SK-N-SH, significant proband enrichment was observed for 3′ UTR variants (*p*_*diff*_ = 19.54%; p = 0.0056), missense variants (*p*_*diff*_ = 13.48%; p = 0.0442), and splice polypyrimidine tract variants (*p*_*diff*_ = 100%; p = 0.0072).

**Table 1.**
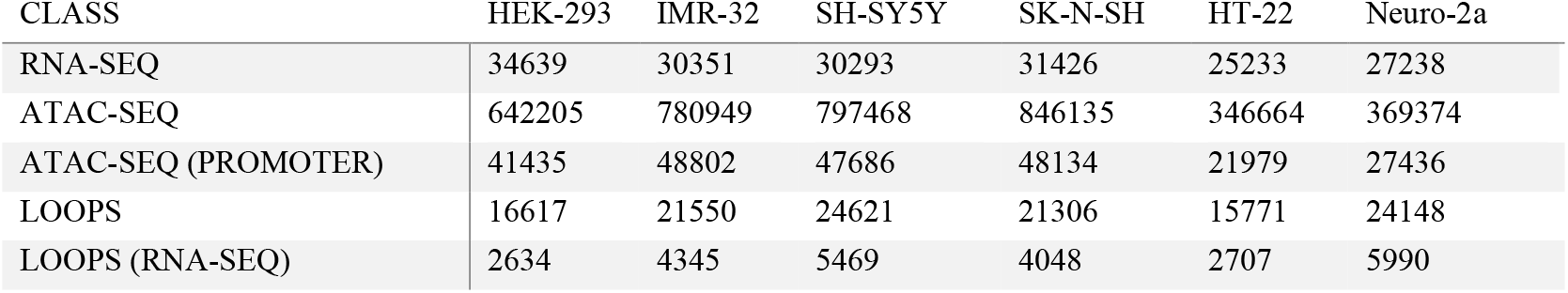
Summary Counts of functional data within cell line for each category. RNA-seq expression data in genes, open chromatin ATAC-seq regions, ATAC-seq with peaks hitting intersecting a promoter region, Hi-C loops, and Hi-C loops overlapping a promoter with RNA expression

| CLASS | HEK-293 | IMR-32 | SH-SY5Y | SK-N-SH | HT-22 | Neuro-2a |
| --- | --- | --- | --- | --- | --- | --- |
| RNA-SEQ | 34639 | 30351 | 30293 | 31426 | 25233 | 27238 |
| ATAC-SEQ | 642205 | 780949 | 797468 | 846135 | 346664 | 369374 |
| ATAC-SEQ (PROMOTER) | 41435 | 48802 | 47686 | 48134 | 21979 | 27436 |
| LOOPS | 16617 | 21550 | 24621 | 21306 | 15771 | 24148 |
| LOOPS (RNA-SEQ) | 2634 | 4345 | 5469 | 4048 | 2707 | 5990 |

**Table 2.** Functional annotations for DNVs with a higher percentage of regions with high PIP in probands versus control siblings. The table displays the percentage of variants in probands and siblings located within regions with high PIP scores, the difference in percentages between the two groups, the odds ratio indicating the likelihood of a variant being in these regions for probands compared to siblings, and the p-values from both Fisher’s exact test and the proportion test.

| Cell Line | Consequence | Total |  | MoDIFI top 5% |  | Diff | Proportion test |
| --- | --- | --- | --- | --- | --- | --- | --- |
|  |  | Proband | Sib | Proband | Sib |  |  |
| IMR-32 | Downstream gene variant | 82 | 75 | 21.95% | 8.00% | 13.95% | 0.0076 |
|  | Missense variant | 85 | 57 | 25.88% | 12.28% | 13.60% | 0.0244 |
| SH-SY5Y | Intergenic variant | 250 | 208 | 11.20% | 6.73% | 4.47% | 0.0495 |
|  | Splice polypyrimidine tract variant | 6 | 7 | 33.33% | 0.00% | 33.33% | 0.0484 |
| SK-N-SH | 3 prime UTR variant | 66 | 58 | 33.33% | 13.79% | 19.54% | 0.0056 |
|  | Missense variant | 74 | 41 | 25.68% | 12.20% | 13.48% | 0.0442 |
|  | Splice polypyrimidine tract variant | 1 | 5 | 100.00% | 0.00% | 100.00% | 0.0072 |

## DISCUSSION

Our work demonstrates that integrating multi-omics data across key neuronal and other cell lines within our probabilistic framework, MoDIFI, can reveal differences across cellular contexts while generating regulatory maps that capture differential activity and mechanistic patterns. This was demonstrated across multiple contexts, including diverse neuronal cell lines, psychiatric traits, regional annotations, and variant classes, underscoring the flexibility and broad applicability of the framework.

First, when examining HGNC supergene family enrichment across all neuronal models, synaptic-related functions emerged as a common functional theme, although mediated by distinct gene families in different cell lines. IMR-32 emphasized synaptic programs associated with early neurodevelopment, including epigenetic regulation and membrane organization; SH-SY5Y highlighted receptor-mediated signaling and plasticity; and SK-N-SH reflected synaptic organization and trafficking consistent with more mature maintenance processes. These results suggest that synaptic regulation represents a convergent functional axis that is differentially deployed across neuronal models, reflecting distinct cellular states rather than a single developmental stage. Finally, because MoDIFI integrates Hi-C, ATAC-seq, and RNA-seq data, it preferentially prioritizes genes supported by coordinated three-dimensional chromatin contacts, regulatory accessibility, and transcriptional output. Such multi-omics concordance is characteristic of neuronal and synaptic regulatory programs and likely contributes to the prominence of synapse-related functional themes observed in the enrichment results.

Next, looking across PGC variants and traits, significant trends between multi-omics PIP and GWAS association strength were generally consistent with those observed for corresponding latent factors derived from cross-disorder analyses. For example, enrichment patterns for eating disorders mirrored those for the Compulsive Disorders factor (F1), both showing strong neuronal cell line effects. Similarly, associations for major depressive disorder aligned with the Internalizing Disorders factor (F4), albeit with more modest trends. Schizophrenia and bipolar disorder exhibited parallel patterns to the Schizophrenia/Bipolar factor (F2), with notable enrichment in HEK-293 and neuronal lines, while ADHD and related neurodevelopmental traits showed correspondence with F3. Substance use disorders demonstrated strong concordance with F5, reinforcing mechanistic overlap across these dimensions. These consistencies suggest that multi-omics integration captures biologically meaningful signals that extend across both individual phenotypes and broader latent dimensions. More broadly, the observed significant trends and cell line-specific differences across PGC traits highlight the potential of MoDIFI to distinguish context-dependent functional activity while prioritizing regions of targeted biological relevance.

Finally, in analyses focused on ASD, signatures of multi-omics neuronal cell line differentiation were evident across multiple levels, including gene set enrichment, implicated enhancers, GWAS variants, and sequence context comparisons between DNVs in ASD probands versus control siblings. This shows the potential utility of MoDIFI in determining the optimal cell line to study the effect of regulatory noncoding variants in different neurodevelopmental disorders. IMR-32 showed elevated MoDIFI levels in the ASD associated gene set, and further tests across the PGC-ASD cohort demonstrated significant GWAS enrichment patterns consistent with those observed for other neurodevelopmental and related traits. Notably, trends in SH-SY5Y mirrored those seen for eating disorders and schizophrenia, as well as latent factors F1 (Compulsive Disorders) and F2 (Schizophrenia/Bipolar), underscoring shared biological mechanisms and convergence between individual trait associations and broader cross-disorder dimensions. While analysis showed consistent trends in MoDIFI PIP levels for enhancer elements, particularly across neuronal cell lines for CNEs, and for HARs in SH‐SY5Y, consistent with its broad neuronal enhancer activity and responsiveness across diverse neurodevelopmental regulatory contexts. Furthermore, ASD proband-specific DNVs exhibited high multi-omics PIP in both coding and noncoding contexts.

Although our model shows promising results in different contexts, future directions are still worth exploring and expanding of the model. From an experimental data perspective, there were a limited number of cell lines for which we generated multi-omics data. Furthermore, single cell lines can be a mixture of different cell types and in those cases single-cell-based multi-omic strategies would be helpful. From a modeling perspective, although specific attention^20^ (see Methods and **Error! Reference source not found**. and **Error! Reference source not found**.) was taken to directly control for inference in combining cross modularity multi-omics data, merging the joint inference could be further investigated and likely improved. Not all cell lines have the same relative functionality, and therefore creating an expanded multilevel Bayesian model explicitly modeling group structure (e.g., neuronal, metabolic, immune) would allow for more refined and nuanced inference. Empirical Bayes priors could leverage extended information in the future to further refine inference.

In summary, our work demonstrates that integrating multi-omics data across key neuronal and other cell lines within the MoDIFI probabilistic framework can reveal functional differences across cellular contexts while generating regulatory maps that capture differential activity and mechanistic patterns. This was shown across diverse contexts, including neuronal cell lines, psychiatric traits, regional annotations, and variant classes, underscoring the flexibility and broad applicability of the approach. These maps provide insight into both shared and context-dependent biology and highlight trends with association statistics that could help strengthen borderline genetic signals. Together, the MoDIFI method and software, along with the accompanying cell‐line annotations, processed ATAC‐seq and RNA‐seq datasets, and other summary‐level resources, create a foundation for guiding future targeted experimentation toward the most relevant cell lines and their paired regulatory or genic regions, ultimately advancing efforts to connect noncoding variation to functional impact.

## METHODS

### Experimental Approach and Biological Materials Cell Lines and Culture Conditions

Cell lines were acquired in accordance with Washington University in St. Louis policies. Following consultation with the Human Research Protection Office, we were informed that sequencing data from the human cell lines cannot be shared. However, summary-level information can be shared upon reasonable request, including aggregate quality-control metrics and non-identifying, gene- or region-level results that do not enable re-identification.

Six cell lines were profiled: the non-neuronal human cell line HEK-293; the human neuronal cell lines IMR-32, SH-SY5Y, and SK-N-SH; and the mouse neuronal cell lines Neuro-2a and HT-22. Cell lines were recently acquired from ATCC or Sigma and cultured under standard conditions (37°C, 5% CO_2_). Media were as follows: HEK-293, DMEM (high glucose) + 10% fetal bovine serum (FBS) + Penicillin-Streptomycin (Pen-Strep) + L-Glutamine; IMR-32 and SK-N-SH, EMEM (high glucose) + 10% FBS + Pen-Strep + L-Glutamine; SH-SY5Y, EMEM (high glucose) + 20% FBS + Pen-Strep + L-Glutamine.

### Karyotyping

Cells were expanded in T25 flasks under standard conditions and harvested at 60-70% confluency. Samples were processed by the Washington University in St. Louis Cytogenetics and Molecular Pathology Core Laboratory. Cells were treated with hypotonic solution, fixed, and stained using Giemsa (GTG) banding. Twenty metaphases per cell line were examined using microscopy and CytoVision imaging software.

### Illumina Short-Read WGS

Genomic DNA was extracted from ∼3 million cells using the Maxwell RSC Cultured Cells DNA Kit. Libraries were sequenced on an Illumina NovaSeq 6000 at the McDonnell Genome Institute (Washington University in St. Louis). FASTQs were aligned to mm10 using bwa mem (v0.7.17-r1188;),^56^ followed by sorting and indexing with SAMtools (v1.9).^57^ Coverage depth was computed per-base and in 5,000 bp windows using mosdepth (v0.3.3).^58^ SNVs/indels were called using DeepVariant (v1.0.0).^59^ Structural variants were called using manta (v1.6.0).^60^

### PacBio HiFi Long-Read WGS

High-molecular weight (HMW) DNA was extracted using the Quick-DNA HMW MagBead Kit from ∼3 million cells per sample; 6–8 samples were pooled after extraction to obtain sufficient input material. Manufacturer protocols were followed with the following modifications per sample: Liquid Sample Buffer 300 µL, Biofluid & Solid Tissue Buffer 300 µL, Quick-DNA MagBinding Buffer 600 µL, MagBinding Beads 100 µL, and DNA Elution Buffer 150 µL. Samples were initially mixed 3 to 4 times using standard pipette tips to break clumps, then mixed using wide-bore tips to minimize shearing. Pooled DNA was quantified using the Qubit dsDNA Broad Range Assay and stored at 4°C to minimize freeze-thaw cycles. PacBio HiFi WGS was performed either on a PacBio Sequel II (McDonnell Genome Institute, Washington University in St. Louis) or on a PacBio Revio (Maryland Genomics, Institute for Genome Sciences, University of Maryland School of Medicine).

Two analysis strategies were used: 1) *de novo* assembly. Assemblies were generated using HiCanu (v2.2)^61^ and Hifiasm (v0.16.1-r375)^62^ reference-based analysis. Reads were mapped to mm10 using pbmm2 (v1.10.0; https://github.com/PacificBiosciences/pbmm2), then merged/sorted/indexed using SAMtools (v1.6). SNVs/indels were called using DeepVariant via NVIDIA Parabricks (v4.1.0-1).^63^ Structural variants were called with pbsv (v2.9.0; https://github.com/PacificBiosciences/pbsv). SVs were merged and compared using SURVIVOR (v1.0.7)^64^ with parameters: maximum breakpoint distance 10 bp; minimum SV size 50 bp; SV type and strand considered; and genComp used for comparison. 5mC detection. CpG methylation (5mC) was inferred using pb-CpG-tools (aligned_bam_to_cpg_scores v2.3.1; https://github.com/PacificBiosciences/pb-CpG-tools) with pileup_calling_model.v1.tflite, minimum mapq 30, and minimum coverage 10.

### ATAC-Seq

ATAC-seq libraries were generated from 1×10^5^ cells using the Active Motif ATAC-Seq Kit following the manufacturer protocol with a modified SPRI cleanup: 35 µL beads were added to the 50 µL sample (0.7×), supernatant retained, then 93.5 µL beads were added (1.1×); beads were washed and DNA eluted to complete the protocol. Unique index primer pairs were used for each replicate. Libraries were sequenced on an Illumina NovaSeq 6000 at the McDonnell Genome Institute (Washington University in St. Louis), targeting 100 million paired-end read pairs per replicate. FASTQs were processed using the ENCODE ATAC-seq pipeline (https://github.com/ENCODE-DCC/atac-seq-pipeline)^65^

### Hi-C

Hi-C libraries were prepared using the Arima Hi-C Kit and Arima Hi-C Library Kit following manufacturer protocols with unique index primer pairs per replicate. Libraries were sequenced on an Illumina NovaSeq 6000 at the McDonnell Genome Institute (Washington University in St. Louis), targeting 1 billion paired-end read pairs per replicate. FASTQs were processed using the ENCODE Hi-C uniform processing pipeline (https://github.com/ENCODE-DCC/hic-pipeline). ^65^

### RNA-seq

RNA-seq libraries were prepared from 1.8×10^6^ cells using the Maxwell RSC SimplyRNA Cells Kit per the manufacturer protocol. Libraries were sequenced on an Illumina NovaSeq 6000 at the McDonnell Genome Institute (Washington University in St. Louis), targeting 200 million paired-end read pairs per replicate. FASTQs were processed with the ENCODE RNA-seq pipeline (https://github.com/ENCODE-DCC/rna-seq-pipeline). ^65^

### PacBio Iso-Seq

RNA was extracted using the Maxwell RSC SimplyRNA Cells Kit. Three replicate RNA preparations were generated, and the sample with the highest RIN was selected for Iso-Seq. Iso-Seq was performed on a PacBio Sequel II at the McDonnell Genome Institute (Washington University in St. Louis). HiFi reads were processed using lima (https://github.com/PacificBiosciences/barcoding), isoseq3 refine, isoseq3 cluster, pbmm2 alignment to mm10.fa, isoseq3 collapse, and pigeon (REF: https://github.com/PacificBiosciences/pigeon) to sort, index, classify, filter, and report isoforms and annotations using mm10.fa and GENCODE vM31.^66^

### Data Processing, Feature Construction, and Multi-Omic Linkage

#### Cross-Species Gene Processing and Superfamily Definitions

For cross-species comparisons involving the two mouse cell lines, orthologous genes were linked between mouse and human. GENCODE annotations were used to establish genomic coordinates, and Ensembl canonical transcripts were used based on human hg38 and mouse mm10 reference genomes. Liftover^67,68^ was used to obtain human genomic coordinates from mouse where needed. Gene sets were restricted to superfamilies containing at least 10 genes.

#### Differential Analysis of ATAC-seq and RNA-seq

ATAC-seq, RNA-seq, and Hi-C data were generated with three biological replicates per cell line. Gene annotations were obtained from the hg38 referqence genome and used for downstream annotation and integrative analyses. For ATAC-seq, high-confidence peak regions were identified and pooled across all cell lines using BEDTools^69^ to define a consistent set of genomic intervals for comparative analysis. Read counts were quantified for each pooled ATAC-seq region across all replicates to generate a region-by-sample count matrix. Low-confidence regions were filtered by removing peaks with quality scores Q < 5. RNA-seq reads were quantified at the gene level, and expected counts were used to build gene-by-sample count matrices and corresponding sample annotation tables. Differential chromatin accessibility and differential gene expression were performed using DESeq2^19^. For both assays, counts were modeled with a negative binomial distribution using a design formula specifying cell line as the primary factor. Unless otherwise specified, DESeq2 defaults were used for normalization, dispersion estimation, and Wald testing, with Benjamini-Hochberg multiple testing correction. Features producing log2 fold-change estimates with zero p-values (e.g., due to absence of counts across all cell lines) were excluded from downstream analyses.

#### Hi-C loop Definition and Loop-Peak-Gene mapping

Hi-C interaction loops identified using HICCUPs^70^ were used as the structural scaffold for multi-omic integration. Gene annotations were based on GENCODE release 44, and only Ensembl canonical transcripts annotated as protein-coding were used to define promoter gene candidates. ATAC-seq regions overlapping at least 50% of either anchor of a Hi-C loop were identified. Promoters were defined as +500/−1500 bp relative to the transcription start site (TSS). ATAC-seq peaks overlapping at least 50% of the promoter window were linked to candidate genes, and promoter-linked ATAC peaks were associated with loop anchors. This procedure allowed each Hi-C loop to map to one or more promoter-regulatory candidate configurations. For loops associated with multiple candidate combinations, the maximum MoDIFI score was retained to represent the loop, corresponding to the strongest-supported regulatory interaction.

#### Gene and Regulatory Region Annotations

Genes implicated in neurodevelopmental disorders were collected from the literature, including neurodevelopmental disorder genes identified based on rare DNVs,^2,71^ clinically relevant genes from ACMG,^72^ and cross-disorder common variant results from the Psychiatric Genomics Consortium (PGC).^73^ Noncoding regulatory regions were obtained from VISTA^49^ and EnhancerAtlas 2.0.^74^

#### DNV Processing

DNVs were obtained from a prior study.^75^ Variants with allele frequency (AF) ≥ 0.05 in gnomAD^76^ were removed, and alleles observed in both probands and siblings were excluded. Variants were annotated using VEP consequence terms,^77^ and variants lacking an assigned VEP consequence were discarded from analyses. For DNVs, Hi-C loops supported by multiple ATAC-seq and RNA-seq combinations were summarized using the maximum MoDIFI score. When a DNV mapped to multiple Hi-C loops, the mean MoDIFI score across those loops was used to represent the variant.

### Statistical Methods

#### Multi-omics Differential Inference Framework: Integrating Cross Cell Line Expression, Chromatin Activity, and Interaction

We developed MoDIFI, a Bayesian framework to integrate cross cell-line evidence from gene expression, chromatin accessibility, and 3D genome interaction. The objective is to construct a test that provides a quantitative metric for relative differential gene-loop activity in a target cell line compared to all others or a designated baseline (here, HEK-293 serves as the baseline for neuronal cell lines). The workflow proceeds as follows i. calculate summary statistics and perform adjustment for bias and inflation, ii. combine RNA and ATAC Bayes factors within a cell line, iii Combine evidence across cell lines and iv. Calculate posterior inclusion probabilities (PIP). The key output of this framework is the (PIP) for each gene-loop pair, representing the probability that the interaction is functionally differential given the observed data and prior biological knowledge.

First, let observed data, *D* = (*D*_RNA_, *D*_ATAC_), denote the summary statistics derived from DESeq2 for RNA-seq (differential expression) and ATAC-seq (differential accessibility), after bias and inflation control (mixture model).^20^ For each loop-gene pair (*l, g*) in a focal cell line *i*, our target hypothesis *H*_1_: RNA and ATAC both differential, versus *H*_0_: at least one component is not differential. The evidence for *H*_1_ relative to *H*_0_ given the observed data *D* is summarized by the BF:

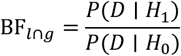

The BF_*l*∩*g*_ is estimated from summary statistics, where we approximate the joint BF by component BFs for RNA and ATAC. Specifically, ATAC evidence partitioned into two interpretable subcomponents for the promoter end of Hi-C loop, BF^Pro^, and the other potentially regulatory end of Hi-C loop BF^Reg^ and then a contribution from the BF^RNA^ differentially RNA expression as well. Under the working assumption of conditional independence under the null between RNA and ATAC summary z-scores, we combine evidence via a joint BF. For loop–gene pairs (*l, g*) for a given cell line *i* versus at least one other cell line, *j*, the joint *BF*_*l*∩*g*_ is then defined as:

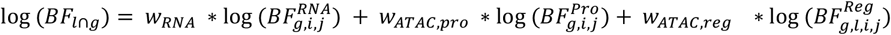

The sum of the weights is constrained such that *Σ w*_*k*_ = 1, where here they were set to equally weight RNA and ATAC-seq evidence *w*_RNA_ = 0.5, *w*_ATAC,pro_ = 0.25, *w*_ATAC,reg_ = 0.25. More generally, this could be expanded to K different experimental or functional lines of evidence, 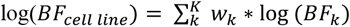. This formulation can be thought of as a weighted geometric mean of Bayes factors^78^ and the same principle is applied later when combining evidence across multiple baseline cell lines.

The BF are approximated utilizing Johnson’s method ^79,80^ to obtain a test‐based Bayes Factor^79–83^ similar to Wakefield’s Bayes factor ^84,85^ (as noted by Wakefield^84^) but more appropriate for generalized linear models. The z value is taken from DESeq2 output, which is based on fitting a negative binomial model. Prior to using summary statistics, we control for bias and inflation with a mixture model^20^. This step is critical to improve interpretability of cross‐functional modularity (here, RNA‐seq versus ATAC‐seq) dramatically reducing inflation (**Error! Reference source not found**. and **Error! Reference source not found**.). BFs are computed separately for RNA‐seq differential expression and ATAC‐seq differential accessibility from the adjusted summary statistics (calculation detailed below under Test‐Based One‐Sided Bayes Factor for GLM Models). When using multiple cell lines as a baseline comparator (as is the case with HEK-293) a weighted average of *log*_10_(*BF*_*meta*_) across C cell lines is calculated:

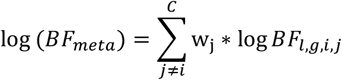

Without prior information, the weighting is evenly distributed across cell lines: 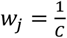, so for HEK-293 this correspond to a weighting of 1/3. For the neuronal cell lines, only HEK-293 was used as a baseline to avoid diluting similar related cell line experimental results.

The *BF*_*meta*_ and empirical priors are then used to calculate the posterior inclusion probabilities (PIP):

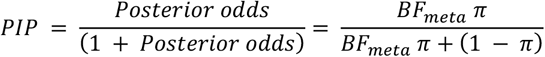

We adopt an empirical Bayes strategy to inform priors *π*_(*l,g,i*)_ by leveraging orthogonal evidence of evolutionary constraint. Intolerance and constraint measures^76,86–92^ are highly effective for identifying genomic regions under negative selection and are widely used to prioritize diagnostic studies.^1,2,93^ Specifically, priors are set proportional to constraint scores derived from Hi-C loop regions, such that *π*_(*l,g,i*)_ ∝ avg(Gnocchi loop regions), where the Gnocchi^21^ score reflects negative selection, computed by transforming the Gnocchi scan statistics to their corresponding percentile to then use as priors.

Higher *π* values correspond to more constrained regions, which are hypothesized to be functionally important. This approach provides biologically motivated prior information that complements RNA and ATAC evidence.

#### Test Based One Sided Bayes Factor for GLM Models

The BF were calculated based on summary statistics after controlling for bias and inflation, assuming that there are g-priors on the corresponding coefficient:

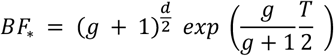

In the model d corresponds to the degrees of freedom, g is a factor based on the g-priors^94^ and T corresponds to a test statistic, say *z*^2^ reported from a model, like results from DESeq2. The corresponding g value is approximated using the empirical Bayes approach as detailed by Held^82,83^:

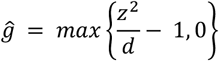

The Bayes factors can be used as a flexible way to model the combined effects across experimental data; however, because we are interested in the one-sided test, if there is significant increased fold change in the cell line being tested (where we are not interested is decrease), the BF needs to be reframed to take this into account. Let’s say that *M*_0_ corresponds to a baseline model where the coefficient we’re making inference on is zero *H*_0_: *β* = 0, then *M*_1_ corresponds to the one sided alternative *H*_1_: *β* > 0 and *M*_2_ corresponds to the two-sided alternative *H*_2_: *β* ≠ 0. Here, *M*_0_ is nested within *M*_1_, which in turn is nested within *M*_2_. This nesting allows us to express the one-sided BF relative to the baselines model as^95^:

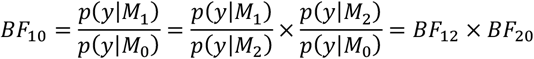

We then use the encompassing prior method of Klugkist et al. such that the one-sided model prior is a truncation distribution of the prior under the two-sided model.^96,97^

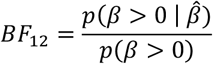

We assume *BF*_20_ is that of the described earlier using g-priors, 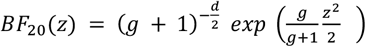, where moving forward we’ll assume d=1 as is the case here. Here *T* = *z*^2^ Wald statistic and g is the scaling factor for the g prior *β*∼*N*(0, *g*(*X*^*T*^*X*)^−1^) or for simplicicty let’s say *β*∼*N*(0, *gV*) for model 2 and *β*∼*N*^+^(0, *gV*) which is the truncated normal for model 1. The posterior has been shown to be^80,83,98^:

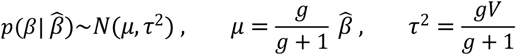

So then plugging in and solving knowing that our prior *p*(*β* > 0)=0.5 since we assume it’s a normal distribution centered at 0:

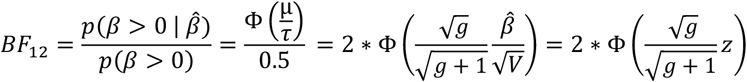

So, the final one-sided Bayes factor is then

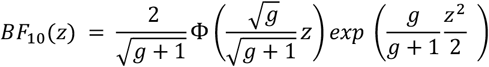

#### Detailed Derivation of the Klugkist Encompassing Prior Bayes Factor

We aim to compute the BF comparing *H*_1_: *θ* > 0 to the encompassing model *H*_0_: *θ* = 0 to show that:

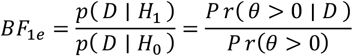

First looking at the encompassing model, we have:

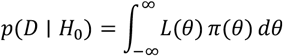

In the encompassing prior approach, we do not specify a separate prior for *H*_1_. Instead, we truncate the prior *π*(*θ*) over the restricted region *R* = (0, ∞), and normalize it:

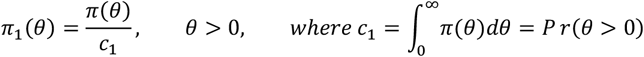

Realizing that *c*_1_ is a constant corresponding to the denominator term we are looking to derive. Then the marginal likelihood under *H*_1_ is:

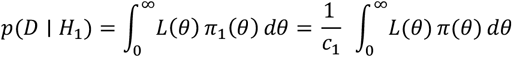

Let’s plug back into the BF.

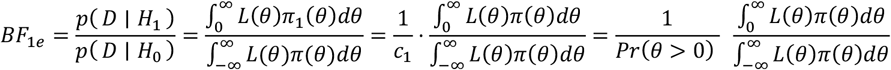

That just leaves the numerator, the posterior probability that *θ* > 0 is defined as.

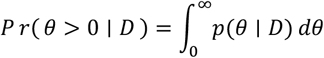

So, the posterior is then defined as:

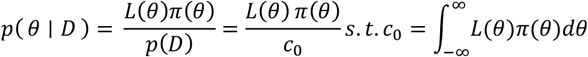

So now notice:

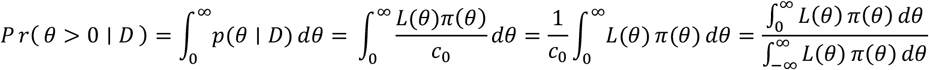

Which can simply be plugged back into the above to show the end equation of interest.

#### MoDIFI Pipeline Implementation

MoDIFI was implemented in Nextflow DSL2. The workflow requires Nextflow (≥23.10),^99^ Java (≥11), and a container runtime (Docker or Singularity/Apptainer). Inputs include promoter annotations (Promoter.tsv), ATAC-seq peak files (.narrowPeak) and alignments (.bam), RNA-seq gene quantification files (*.genes.results), and Hi-C data provided via a resources directory. MoDIFI generates count matrices, annotation tables, and sample condition files for DESeq2 (*counts.txt, *ann.txt, *conds.txt), and supports re-analysis using customized target-reference comparisons without rerunning upstream preprocessing.

### External Datasets and Enrichment Analyses

#### Psychiatric Genomics Consortium GWAS mapping

GWAS summary statistics were obtained from the Psychiatric Genomics Consortium (PGC). Association strength was represented by -log_10_(P). Variants originally mapped to hg19 were lifted over to hg38 using a Python liftover implementation (get_lifter). Lifted variants were mapped to Hi-C loop anchors to assign MoDIFI scores; when a variant mapped to multiple loops, the mean MoDIFI score was used per variant. Variants were stratified into significance bins using -log_10_(P) thresholds (>2, >3, >4, >5) with -log_10_(P) < 2 as the reference. Genes were categorized as high-confidence (MoDIFI ≥ 0.95) or lowconfidence (MoDIFI < 0.95). Odds ratios were computed to evaluate enrichment of high-MoDIFI genes across GWAS significance thresholds.

#### Gene Family Enrichment Analysis

Gene family enrichment was assessed using a hypergeometric over-representation test. For genes with multiple MoDIFI scores, gene-level values were summarized using the median. Enrichment was evaluated by comparing the proportion of genes within each family with median MoDIFI ≥ 0.95 against expectation under either a global background or a superfamily-specific background. ASD-associated genes were defined using the curated gene list from the PGC ASD 2019 GWAS.^46^

## Supplemental Data

Supplemental data 1 is a table with rows representing genes and columns for each cell line, indicating gene expression along with identifies whether each gene is a known ASD gene. Supplemental data 2 is in BED format and contains all predicted ATAC-seq peaks for each cell line. This table similarly includes columns indicating whether the peak is open or not. Supplemental data 3 delineates all ATAC-seq peaks linked to gene promoters. It includes a column indicating whether each gene is a known ASD gene and additional columns showing if the link exists, with yes/no values for each cell line. Summary results of the data demonstrate small proportion of regions with open chromatin that also hit promoters and Hi-C those loops that correspond to RNA-seq expression data as well. These layered functional evidence and key regions with limited intersection across all regulatory data represent more likely targets for testing for biologic function.

## DECLARATIONS

Ethics approval and consent to participate: not applicable. Consent for publication: not applicable. Availability of data and materials: Mouse cell line data is available at https://www.ncbi.nlm.nih.gov/bioproject/?term=PRJNA938057. We do not have permission to share the underlying sequence data from the human cell lines. Competing interests: none. Funding: This study was funded by the National Institutes of Health (R01MH126933). Author contributions: A.V.R. performed computational analyses, wet-lab experiments, and wrote the manuscript. G.T.S. performed computational analyses, implemented statistical methods and software, and wrote the manuscript. J.M. performed wet-lab experiments. T.L.M. performed computational analyses. H.H. performed wet-lab experiments. J.K.N. performed computational analyses. H.K. performed computational analyses and experiments. T.J.H. conceptualized the study, derived and developed the statistical methods and framework, performed computational analyses, and wrote the manuscript. T.N.T. conceptualized the study, acquired funding, performed computational analyses and wet-lab experiments, and wrote the manuscript. All authors read and approved the final manuscript. Acknowledgements: Thank you to the McDonnell Genome Institute and Maryland Genomics for their sequencing services.

