## Supplemental Tables and Figures for "Multi-omics Differential Inference for Functional Interpretation (MoDIFI): A Statistical Framework to Prioritize Cell Lines for Neurodevelopmental Variants"

| Cell Line | Consequence | Total Proband | Total Sib | In Loop % Proband | In Loop % Sib | Diff | OR | Fishers p-value | Proportion test |
|---|---|---|---|---|---|---|---|---|---|
| *IMR-32* | Stop gained | 139 | 59 | 15.8% | 3.4% | 12.4% | 5.36 | **0.009** | **0.007** |
| | Intron | 87711 | 68893 | 10.3% | 9.9% | 0.4% | 1.04 | **0.01** | **0.009** |
| *SH-SY5Y* | Stop gained | 139 | 59 | 18.0% | 6.8% | 11.2% | 3.02 | **0.029** | **0.021** |
| | Splice region | 198 | 150 | 20.7% | 12.7% | 8.0% | 1.8 | **0.033** | **0.025** |
| | Stop lost | 1 | 5 | 100.0% | 0.0% | 100.0% | - | 0.167 | **0.007** |
| *SK-N-SH* | Splice donor $5^{th}$ base | 26 | 15 | 23.1% | 0.0% | 23.1% | 0 | **0.051** | **0.022** |

***Supplemental Table 1 Functional annotations for DNVs with higher percentage of Hi-C loop calls in probands versus healthy siblings.*** *Each DNV was annotated with VEP consequences and stratified for variants found in either probands or siblings. The variants were linked to Hi-C interaction data to assess the percentage of variants associated with each functional consequence across different cell lines. This allowed for a comparison of differences between probands and siblings. Displayed are the percentage of variants in probands or siblings that are located within a Hi-C loop call, the difference in loop percentages between probands and siblings, the odds ratio indicating the likelihood of a variant being within a loop for probands compared to siblings, the p-value from Fisher's exact test, and p-value from the proportion test, providing additional statistical support for the results.*

| Cell Line | Consequence | Total Proband | Sib | In ATAC % Proband | Sib | Diff | OR | Fishers p-value | Proportion test |
|---|---|---|---|---|---|---|---|---|---|
| *HEK-293* | *TFBS* | 406 | 349 | *52.2%* | *44.7%* | *7.5%* | 1.35 | **0.023** | **0.02** |
| | *Noncoding transcript exon* | 3952 | 2967 | *12.7%* | 11.2% | 1.4% | 1.15 | **0.038** | **0.035** |
| | *Splice donor region* | 42 | 40 | *21.4%* | 7.5% | 13.9% | 3.36 | 0.069 | **0.037** |
| *IMR-32* | *Synonymous* | 725 | 602 | *22.5%* | 18.8% | 3.7% | 1.26 | 0.056 | **0.049** |
| | *Splice acceptor* | 40 | 19 | *15.0%* | 0.0% | 15.0% | - | 0.085 | **0.037** |
| *SH-SY5Y* | *Intergenic* | 42348 | 32895 | *4.9%* | 4.5% | 0.4% | 1.08 | **0.01** | **0.01** |
| *SK-N-SH* | *Start lost* | 4 | 8 | *50.0%* | 0.0% | 50.0% | - | 0.091 | **0.014** |

**Supplemental Table 2 Functional annotations for DNVs with higher percentage of calls within ATAC-seq open chromatin regions in probands versus healthy siblings.** Variants were linked to ATAC-seq open chromatin regions to assess the percentage of variants in these regions in probands compared to siblings, across specific cell lines and functional annotations. The results include the percentage of variants located within an open chromatin region for both probands and siblings, the difference in percentages of open regions between the two groups, the odds ratio indicating the likelihood of a variant being within an open region for probands compared to siblings, and the p-values from both Fisher's exact test and the proportion test.

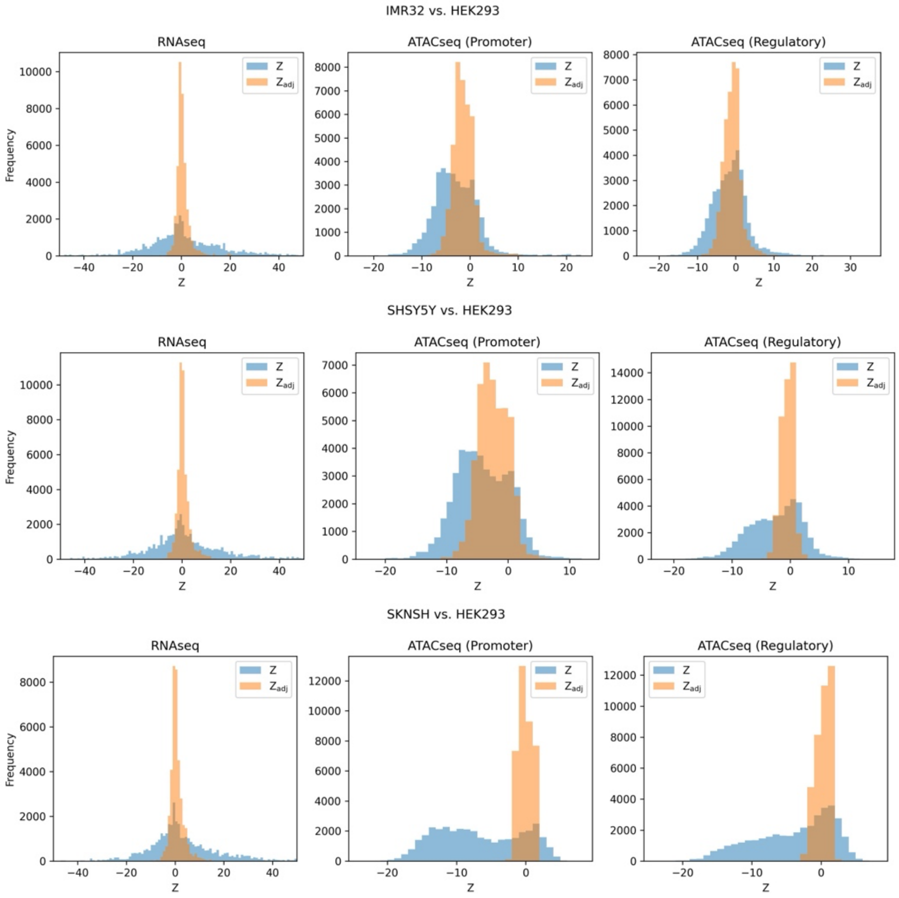


***Supplemental Figure 1 Distribution of Z-scores before and after mixture model adjustment.*** *Histograms show the distributions of original Z-scores (blue) and mixture model adjusted Z-scores (orange) for differential expression analysis of RNA-seq and ATAC-seq (promoter-end and regulatory-end of chromatin loops). Using the mixture model adjustment reduces inflation and variance in the Z-score distributions, resulting in more centered and comparable profiles across data modalities.*

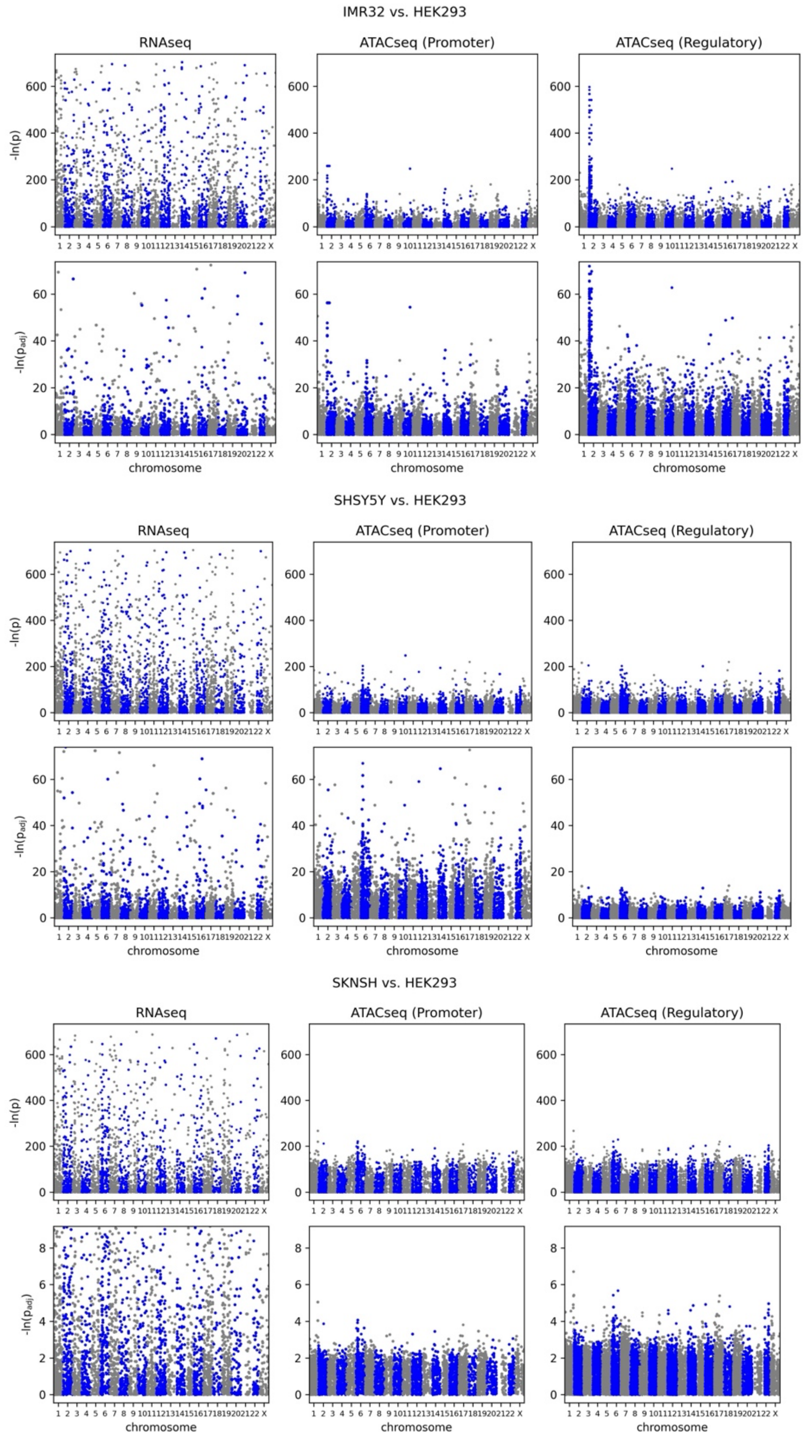


***Supplemental Figure 2 Manhattan plots of differential RNA-seq and ATAC-seq before and after mixture model adjustment.*** *Inflated Z-scores and corresponding p-values from RNA-seq and ATAC-seq (promoter-end and regulatory-end of chromatin loops) were corrected using the* bacon *approach. The y-axis shows* $-ln\ (P)$*. Extreme inflation observed prior to adjustment was substantially reduced for both RNA-seq and ATAC-seq after correction.*

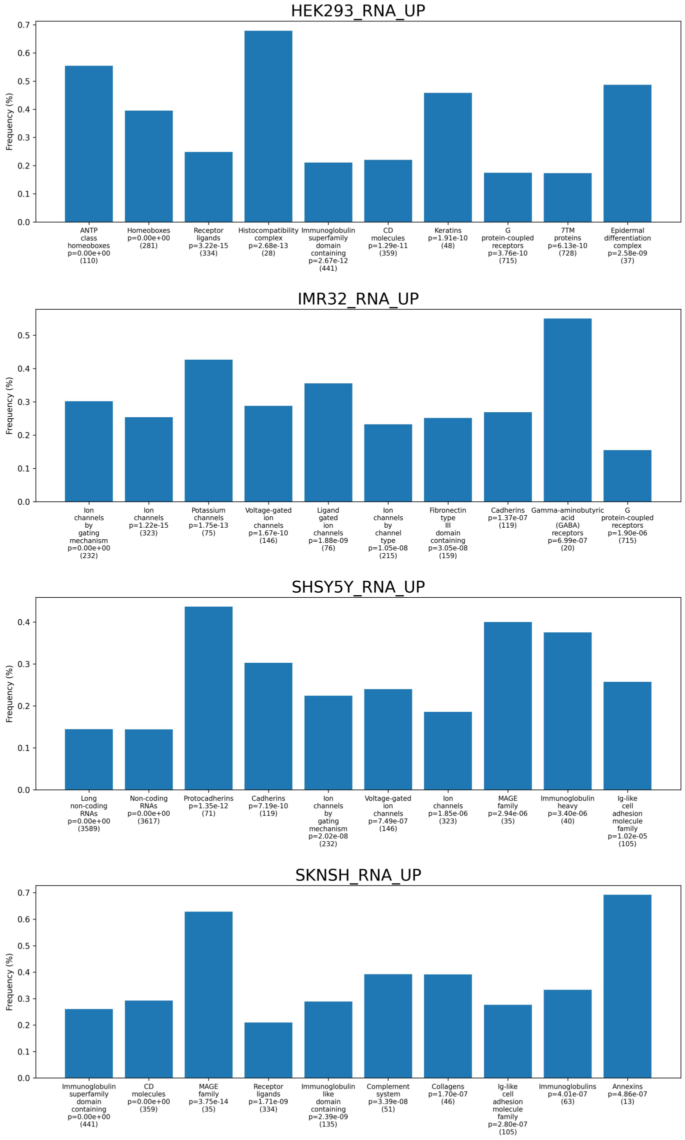


***Supplemental Figure 3* Differential RNA Expression for Super Gene Families in HGNC** *The frequency of genes within each super gene family from HGNC that showed differential expression in the top 10%. We then performed an exact test to determine the probability of observing this level of relative enrichment by chance. The top ten super gene families with elevated frequencies of genes in the top 10% of differentiation for each cell line are displayed. They are sorted by relative significance, with the number of genes in each super gene family indicated in parentheses.*

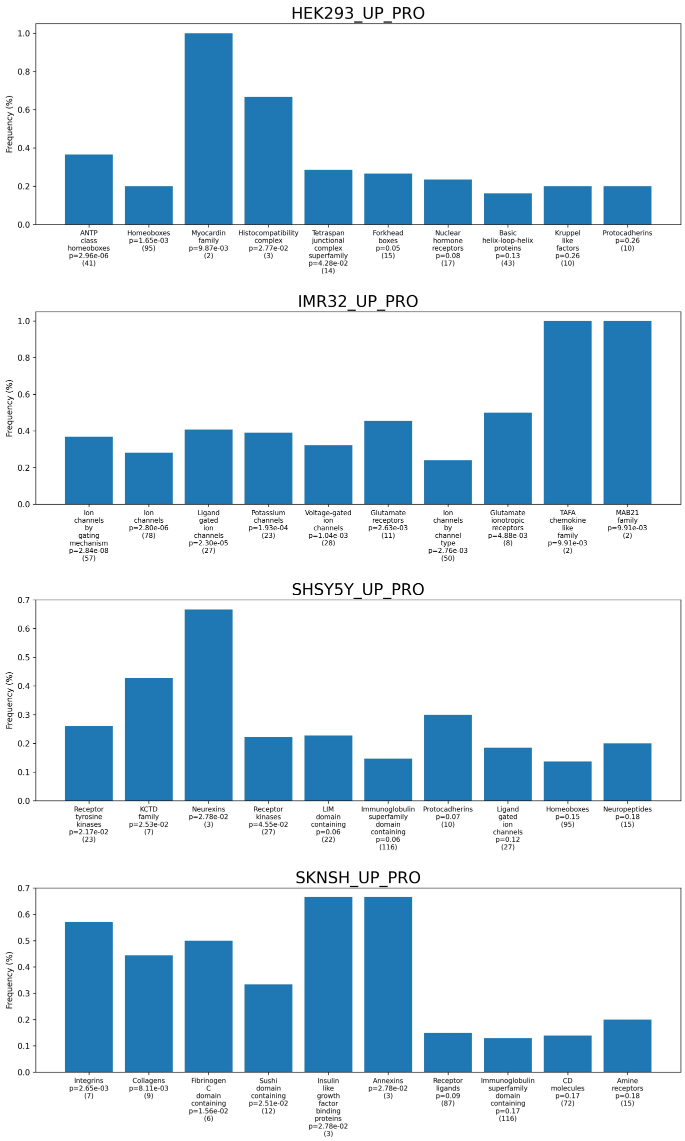


***Supplemental Figure 4 Differential Chromatin Accessibility and Interactions for Super Gene Families in HGNC*** *This figure illustrates the frequency of genes within each super gene family from HGNC that exhibited differential chromatin accessibility and interactions in the top 10%. We conducted an exact test to assess the probability of observing this level of relative enrichment by chance. The top ten super gene families with elevated frequencies of genes in the top 10% for each cell line are displayed, sorted by relative significance, with the number of genes in each super gene family indicated in parentheses.*

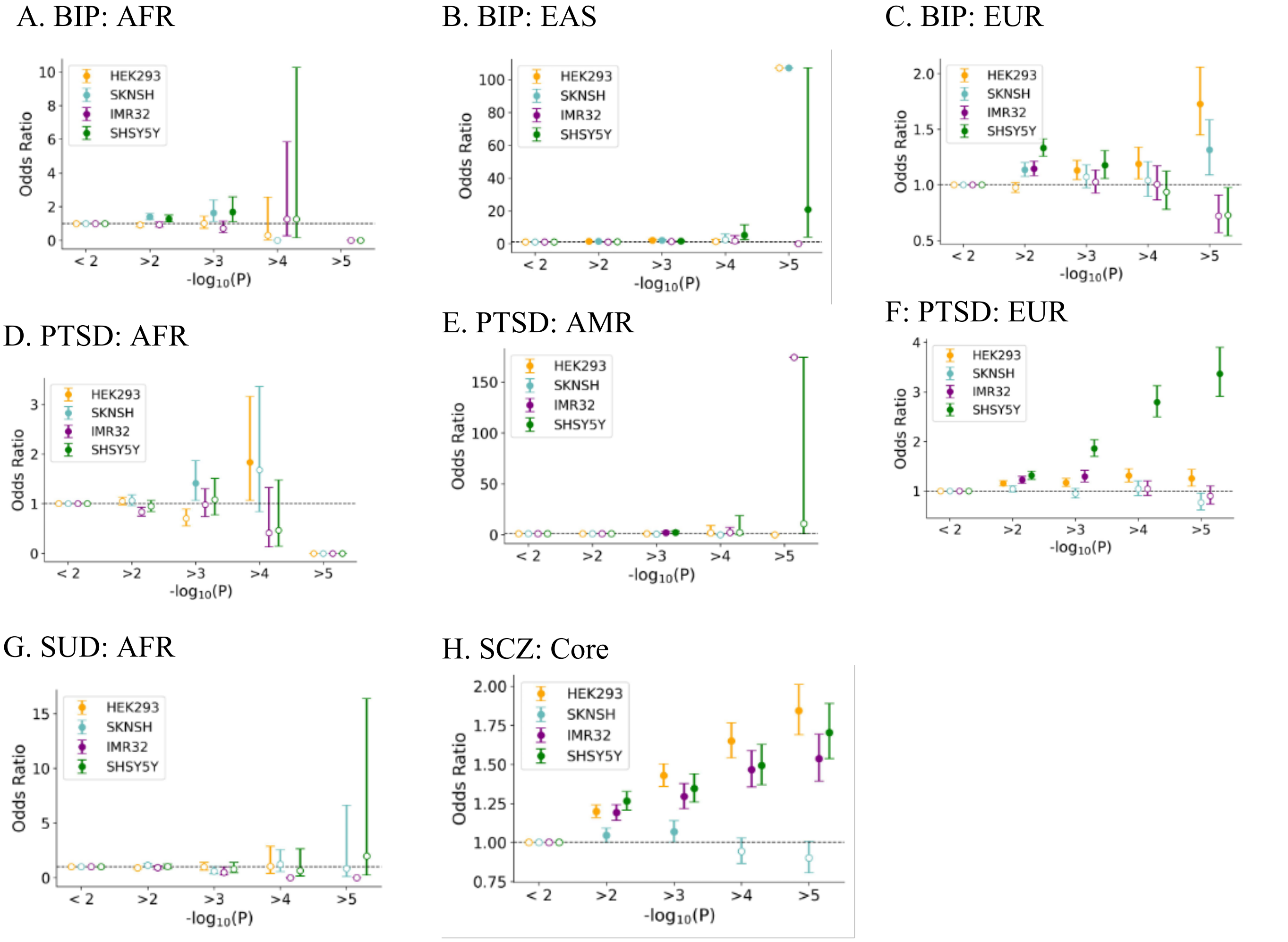


***Supplemental Figure 5** Enrichment of psychiatric traits stratified by genetic ancestry for BIP and PTSD* with *associated variants in high MoDIFI PIP regulatory regions across cell lines*

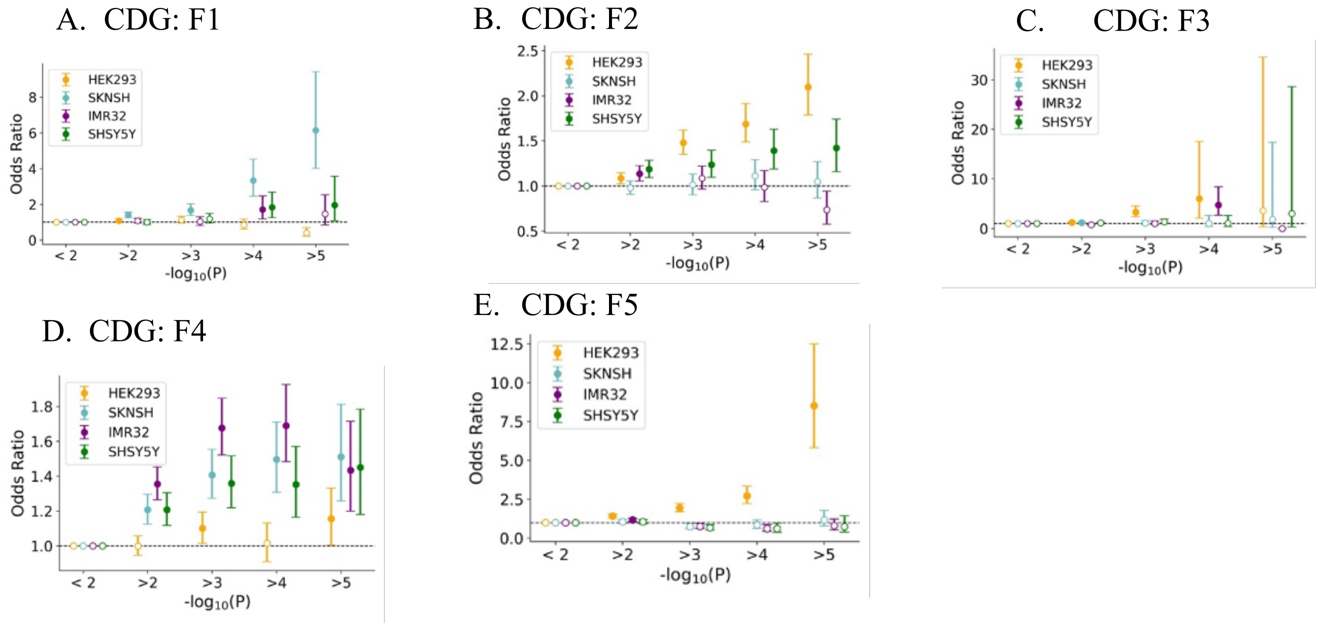


***Supplemental Figure 6 Enrichment of latent factor analysis of psychiatric traits–associated variants in high MoDIFI PIP regulatory regions across cell lines*** *For each PGC trait, panels display the odds ratio of SNPs overlapping the top 95th percentile of posterior inclusion probability (PIP) scores versus those below the 95th percentile across four cell lines (*HEK-293, *SK-N-SH, IMR-32, and SH-SY5Y). SNPs are binned by increasing GWAS significance (−log10(p) thresholds of <2, >2, >3, >4, and >5 on the x-axis), and points represent odds ratios with 95% confidence intervals error bars. Horizontal dashed lines denote no enrichment (OR = 1). For all plots, solid dots indicate significant OR at 5% alpha level whereas hollow ones are not significant.* Panes correspond to latent factor analysis clustered phenotypes corresponding to: *A.) F1, a Compulsive disorders factor defined by AN, OCD and, more weakly, TS and ANX;* B.) *F2, a SB factor defined by SCZ and BIP;* C.) *F3, a Neurodevelopmental factor defined by ASD, ADHD and, more weakly, TS;* D.) *F4, an Internalizing disorders factor defined by PTSD, MD and ANX; and* D.) *F5, a SUD factor defined by OUD, CUD, AUD, NIC and, to a lesser extent, ADHD*.
