## Supplementary material for "Multi-omics Differential Inference for Functional Interpretation (MoDIFI): A Statistical Framework to Prioritize Cell Lines for Neurodevelopmental Variants": Column Dictionary.docx

Table1

| **Column Name** | **Definition** |
| --- | --- |
| GeneID_Human, GeneName_Human, GeneType_Human | Gene ID, Gene Name and Gene Type from RNA-seq outputs for each human cell line |
| GeneID_Mouse, GeneName_Mouse | Gene ID, Gene Name and Gene Type from RNA-seq outputs for each mouse cell line |
| Chrom_Human, Start_Human, End_Human | For each human gene, genomic coordinates are sourced from the GENCODE (hg38) database, where genes are defined by their Ensembl transcripts. |
| Chrom_Mouse, Start_Mouse, End_Mouse | For each mouse gene, genomic coordinates are sourced from the GENCODE (mm10) database, where genes are defined by their Ensembl transcripts.  Additionally, orthologous genes between human and mouse are retrieved from the Ensembl BioMart accessible at <http://www.ensembl.org/biomart/martview>. |
| TPM_HEK293, TPM_IMR32, TPM_SHSY5Y, TPM_SKNSH, TPM_HT22, TPM_neuro2a | TPM (Transcripts Per Million) is utilized to quantify gene expression in RNA-seq data for each cell line. |
| AutismGene | A list of 379 genes associated with autism is available at the provided [link](https://github.com/TNTurnerLab/HT22_genome_epigenome_functional_genomics_project/blob/main/gene_sets/genes_with_genome-wide_significance_for_de_novo_variants_in_NDDs_from_Coe_et_al_2019_Nature_Genetics_and_Kaplanis_et_al_2020_Nature.txt). If a gene is linked to autism, it is indicated with a 'Yes'." |
| deNovo | Indicates the number of de novo variants affecting the gene. |
| deNovo_HitPromoter | Displays the number of de novo variants impacting the upstream 5kb region of the gene, commonly referred to as the promoter region. |
| most_severe_consequence from VEP | For each de novo variant, the most severe consequences were determined using VEP. At the gene level, the most severe consequences associated with all de novo variants related to the gene are displayed. |
| PGC | Average p-values are calculated for all PGC variants within a given gene that have p-values less than 0.01, as listed in iPSYCH-PGC_ASD_Nov2017.txt. |
| numVariantPGC | Within each gene, the number of PGC variants associated with Autism, as listed in iPSYCH-PGC_ASD_Nov2017.txt, with p-values less than 0.01, are counted. |

Table2

| **Column Name** | **Definition** |
| --- | --- |
| Chrom_Human, Start_Human, End_Human | List all genomic regions identified by ATAC-seq in human cell lines, based on the hg38 reference genome, from narrowPeak output files.  For mouse cell lines, human genomic coordinates were obtained using the LiftOver tool with the chain file mm10ToHg38.over.chain.gz, downloaded from <https://genome.ucsc.edu/cgi-bin/hgLiftOver>. |
| Chrom_Mouse, Start_Mouse, End_Mouse | List all genomic regions identified by ATAC-seq in mouse cell lines, based on the mm10 reference genome, from narrowPeak output files. |
| [cell line]_Signal  [cell line]_pvalue  [cell line]_qvalue  1 human Kidney cell line: HEK293  3 human neuronal cell lines: MR32, SHSY5Y SKNSH.  2 mouse neuronal cell lines  HT22, neuro2a | Signals, p values and q values are collected from ATAC-seq outputs |
| GeneID_Human,  GeneName_Human | ATAC-seq regions are overlapped with ‘**protein-coding**’ genes defined by their Ensembl canonical transcripts in the GENCODE (hg38) database. Both Gene ID and Gene Name are recorded for each overlap. |
| GeneID_Mouse, GeneName_Mouse | ATAC-seq regions are overlapped with ‘**protein-coding**’ genes defined by their Ensembl canonical transcripts in the GENCODE (mm10) database. Both Gene ID and Gene Name are recorded for each overlap. |
| OverLapWithinGene | What percentage of the ATAC-seq region falls within the gene? |
| OverLapinPRO | What percentage of the ATAC-seq region falls within the 5kb upstream region of a gene? |
| AutismGene | A list of 379 genes associated with autism is available at the provided [link](https://github.com/TNTurnerLab/HT22_genome_epigenome_functional_genomics_project/blob/main/gene_sets/genes_with_genome-wide_significance_for_de_novo_variants_in_NDDs_from_Coe_et_al_2019_Nature_Genetics_and_Kaplanis_et_al_2020_Nature.txt). If an ATAC-seq region intersects with any of these autism-associated genes, the corresponding human gene names are indicated. |
| deNovo | Indicates the number of de novo variants within ATAC-seq regions. |
| deNovo_HitPromoter | Indicates the number of de novo variants impacting the upstream 5kb region of the gene, commonly referred to as the promoter region, within ATAC-seq regions |
| most_severe_consequence from VEP | For each de novo variant, the most severe consequences were determined using VEP. Within ATAC-seq region, the most severe consequences associated with all de novo variants are displayed. |
| PGC | Average p-values are calculated for all PGC variants within a ATAC-seq region that have p-values less than 0.01, as listed in iPSYCH-PGC_ASD_Nov2017.txt. |
| numVariantPGC | Within each ATAC-seq region, the number of PGC variants associated with Autism, as listed in iPSYCH-PGC_ASD_Nov2017.txt, with p-values less than 0.01, are counted. |

Table 3

All column definitions in Table 3 are the same as those in Table 2. Additionally, Table 3 includes only variants with an OverLapinPRO value of 50% or higher.
